# Continuous Genetic Recording with Self-Targeting CRISPR-Cas in Human Cells

**DOI:** 10.1101/053058

**Authors:** Samuel D. Perli, Cheryl H. Cui, Timothy K. Lu

**Affiliations:** Synthetic Biology Group, MIT Synthetic Biology Center, Massachusetts Institute of Technology, Cambridge, MA 02139, USA.; Research Laboratory of Electronics, Massachusetts Institute of Technology, Cambridge, MA 02139, USA.; Department of Electrical Engineering & Computer Science, Massachusetts Institute of Technology, Cambridge, MA 02139, USA.; Harvard-MIT Division of Health Sciences and Technology, Cambridge, MA 02139, USA.; Department of Biological Engineering, Massachusetts Institute of Technology, Cambridge, MA 02139, USA.

## Abstract

The ability to longitudinally track and record molecular events *in vivo* would provide a unique opportunity to monitor signaling dynamics within cellular niches and to identify critical factors in orchestrating cellular behavior. We present a self-contained analog memory device that enables the recording of molecular stimuli in the form of DNA mutations in human cells. The memory unit consists of a self-targeting guide RNA (stgRNA) cassette that repeatedly directs *Streptococcus pyogenes* Cas9 nuclease activity towards the DNA that encodes the stgRNA, thereby enabling localized, continuous DNA mutagenesis as a function of stgRNA expression. We analyze the temporal sequence evolution dynamics of stgRNAs containing 20, 30 and 40 nucleotide SDSes (Specificity Determining Sequences) and create a population-based recording metric that conveys information about the duration and/or intensity of stgRNA activity. By expressing stgRNAs from engineered, inducible RNA polymerase (RNAP) III promoters, we demonstrate programmable and multiplexed memory storage in human cells triggered by doxycycline and isopropyl β-D-1-thiogalactopyranoside (IPTG). Finally, we show that memory units encoded in human cells implanted in mice are able to record lipopolysaccharide (LPS)-induced acute inflammation over time. This tool, which we call Mammalian Synthetic Cellular Recorder Integrating Biological Events (mSCRIBE), provides a unique strategy for investigating cell biology *in vivo* and *in situ* and may drive further applications that leverage continuous evolution of targeted DNA sequences in mammalian cells.

**One Sentence Summary:** By designing self-targeting guide RNAs that repeatedly direct Cas9 nuclease activity towards their own DNA, we created multiplexed analog memory operators that can record biologically relevant information *in vitro* and *in vivo*, such as the magnitude and duration of exposure to TNAα.

---

Cellular behavior is dynamic, responsive and regulated by the integration of multiple molecular signals. Biological memory devices that can record regulatory events would be useful tools for investigating cellular behavior over the course of a biological process and further our understanding of signaling dynamics within cellular niches. Earlier generations of biological memory devices relied on digital switching between two or multiple quasi-stable states based on active transcription and translation of proteins (*1*–*3*). However, such systems do not maintain their memory after the cells are disruptively harvested. Encoding transient cellular events into genomic DNA memory using DNA recombinases enables the storage of heritable biological information even after gene regulation is disrupted (*4*, *5*). The capacity and scalability of these memory devices are limited by the number of orthogonal regulatory elements (e.g., transcription factors and recombinases) that can reliably function together. Furthermore, because they are restricted to a small number of digital states, they cannot record dynamic (analog) biological information, such as the magnitude or duration of a cellular event. We recently demonstrated a population-based technology for genomically encoded analog memory in *Escherichia coli* based on dynamic genome editing with retrons (*6*). Here, we present Mammalian Synthetic Cellular Recorders Integrating Biological Events (mSCRIBE), an analog memory system that enables the recording of cellular events within human cell populations in the form of DNA mutations. mSCRIBE uses self-targeting guide RNAs (stgRNAs) that direct CRISPR-Cas activity to repeatedly mutagenize the DNA loci that encodes the stgRNAs.

The *S. pyogenes* Cas9 system from the Clustered Regularly-Interspaced Short Palindromic Repeats-associated (CRISPR-Cas) family is an effective genome engineering enzyme that catalyzes double-stranded breaks and generates mutations at DNA loci targeted by a small guide RNA (sgRNA) (*7*–*9*). Normal sgRNAs are comprised of a 20 nucleotide (nt) Specificity Determining Sequence (SDS), which specifies the DNA sequence to be targeted and is immediately followed by a 80 nt scaffold sequence, which associates the sgRNA with Cas9. In addition to sequence homology with the SDS, targeted DNA sequences must possess a Protospacer Adjacent Motif (PAM) (5’-NGG-3’) immediately adjacent to their 3’-end in order to be bound by the Cas9-sgRNA complex and cleaved (*10*). When a double-strand break is introduced in the target DNA locus in the genome, the break is repaired by either homologous recombination (when a repair template is provided) or error-prone non-homologous end joining (NHEJ) DNA repair mechanisms, resulting in mutagenesis of the targeted locus (*8*, *9*). It is important to note that even though the DNA locus encoding a normal sgRNA sequence is perfectly homologous to the sgRNA, it is not targeted by the standard Cas9-sgRNA complex because it does not contain a PAM.

To enable continuous encoding of population-level memory in human cells, we sought to build a modular memory unit that can be repeatedly written to generate new sequences and encode additional information over time. With the standard CRISPR-Cas system, once a genomic DNA target is repaired, resulting in a novel DNA sequence, it is unlikely to be targeted again by the original sgRNA, since the novel DNA sequence and the sgRNA would lack the necessary sequence homology. We hypothesized that if the standard sgRNA architecture could be engineered so that it acted on the same DNA locus from which the sgRNA is transcribed, rather than a separate sequence elsewhere in the genome, this would yield a self-targeting guide RNA (stgRNA) that should repeatedly target and mutagenize the DNA that encodes it. To achieve this, we modified the DNA sequence from which the sgRNA is transcribed to include a 5’-NGG-3’ PAM immediately downstream of the region encoding the SDS such that the resulting PAM-modified stgRNA would direct Cas9 endonuclease activity towards the stgRNA’s own DNA locus. After a double-strand DNA break is introduced in the SDS-encoding region and repaired via the NHEJ repair pathway, the resulting *de novo* mutated stgRNA locus should continue to be transcribed as a mutated version of the original stgRNA and participate in another cycle of self-targeting mutagenesis. Multiple cycles of transcription followed by cleavage and error-prone repair should occur, resulting in a continuous, self-evolving Cas9-stgRNA system (Fig. 1A). We hypothesized that by biologically linking the activity of this system with regulatory events of interest, mSCRIBE can serve as a memory device that records information in the form of DNA mutations.

**Fig. 1.**
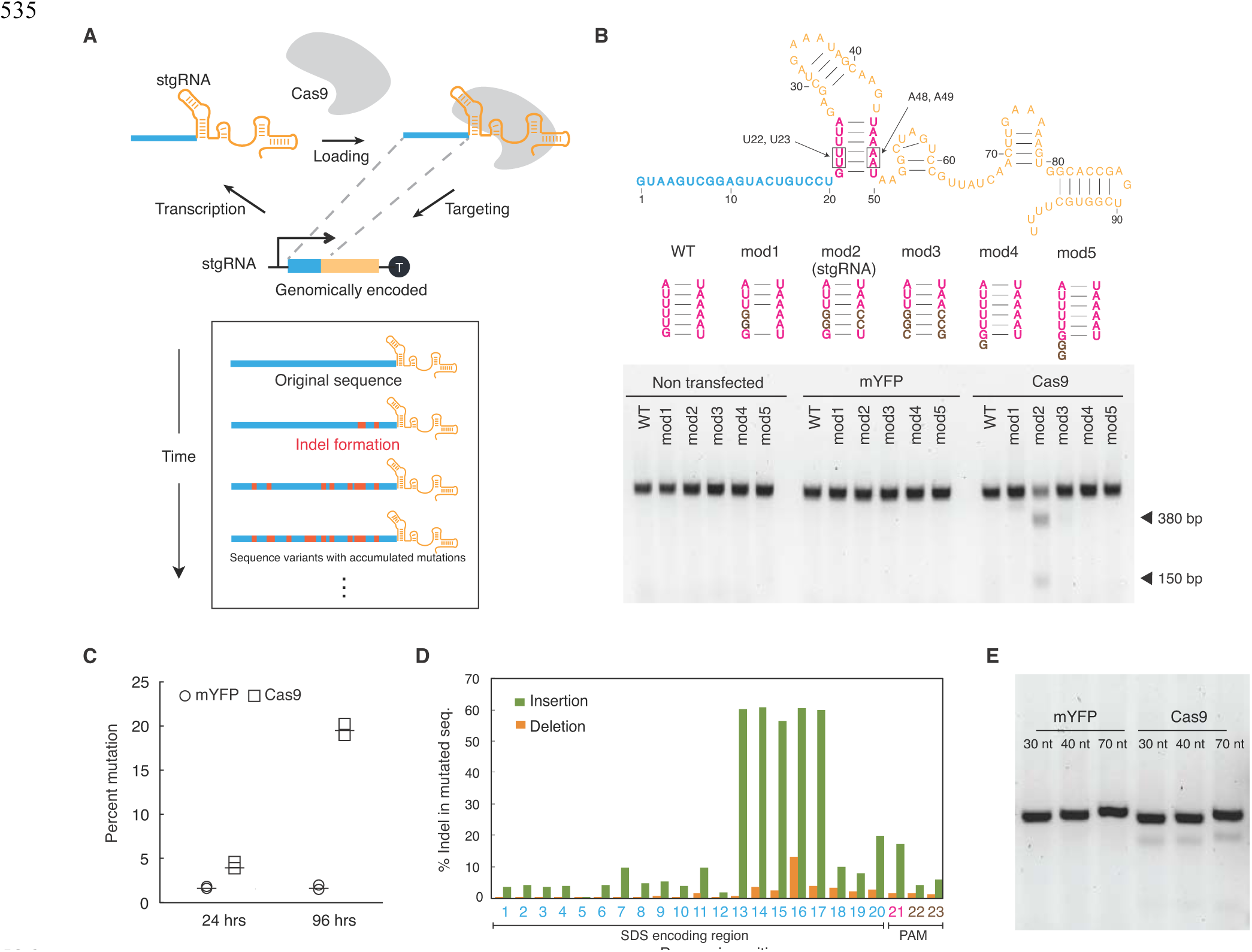
Continuously evolving self-targeting guide RNAs. (**A**) Schematic of the self-targeting CRISPR-Cas system. The Cas9-stgRNA complex cleaves the DNA from which the stgRNA is transcribed, leading to error-prone DNA repair. Multiple rounds of transcription and DNA cleavage can occur, resulting in continuous mutagenesis of the DNA encoding the stgRNA. The blue line in the stgRNA schematic represents the specificity-determining sequence (SDS) while mutations in the stgRNAs are illustrated as red marks. When stgRNA or Cas9 expression is linked to cellular events of interest, accumulation of mutations at the stgRNA locus provides a molecular record of those cellular events. (**B**) Multiple variants of sgRNAs were built and tested for inducing mutations at the DNA loci that encoded them using T7 E1 assays. Introducing a PAM into the DNA encoding the *S. pyogenes* sgRNA (black arrows) renders the sgRNA self-targeting, as evidenced by Cas9-dependent cleavage of PCR amplicons into two fragments (380 bp and 150 bp) in the mod2 sgRNA variant (referred to as stgRNA). HEK 293T derived stable cell lines expressing each of the variant sgRNAs were transfected with plasmids expressing Cas9 or mYFP. Cells were harvested 96 hours post transfection and the genomic DNA was PCR amplified and subjected to T7 E1 assays. (**C**) Further analysis via next-generation sequencing confirmed that stgRNA can effectively generate mutations at its own DNA locus. HEK 293T cells constitutively expressing stgRNA were transfected with plasmids expressing Cas9 or mYFP. PCR-amplified genomic DNA was sequenced via Illumina MiSeq and the percentage of mutated sequences is presented. Only Cas9-transfected cells acquired specific mutations at the stgRNA locus whereas mYFP-transfected cells had a basal level (~1%) mutation rate corresponding to the next-generation sequencing error rate. The data presented above is for two biological duplicates of the experiment. (**D**) The percentage of sequences containing specific mutation types (insertion or deletion) at individual base pair positions out of all mutated sequences is presented. By aligning each of the Illumina MiSeq reads with the original un-mutated stgRNA sequence, the base pair positions of insertions and deletions acquired by the stgRNA locus was calculated. We found higher number of deletions than insertions at each base pair position. (**E**) Computationally designed stgRNAs with longer SDS regions (30nt-1, 40nt-1 and 70nt-1) demonstrate self-targeting activity. HEK 293T-derived stable cell lines expressing 30nt-1, 40nt-1 and 70nt-1 stgRNAs were transfected with plasmids expressing Cas9 or mYFP. T7 E1 assays were performed on the PCR-amplified genomic DNA. Also see Fig. S1 and Constructs 1-11 in Table S2.

## Modifying an sgRNA-expressing DNA locus to include a PAM renders it self targeting

We built multiple variants of a *S. pyogenes* sgRNA-encoding DNA sequence with a 5’-GGG-3’ PAM located immediately downstream of the region encoding the 20 nt SDS, and tested them for their ability to generate mutations at their own DNA locus. HEK 293T-derived stable cell lines were built to express either the wild-type (WT) or each of the variant sgRNAs shown in Fig. 1B (Constructs 1-6, Table S2, Methods). Plasmids encoding either spCas9 (Construct 7, Table S2) or mYFP (negative control) driven by the CMV promoter (CMVp) were transfected into cells stably expressing the depicted sgRNAs, and the sgRNA loci were inspected for mutagenesis using T7 Endonuclease I (T7 E1) assays four days after transfection. A straightforward variant sgRNA (mod1) with guanine substitutions at the U23 and U24 positions did not exhibit any noticeable self-targeting activity. We speculated that this was due to the presence of bulky guanine and adenine residues facing each other in the stem region, resulting in destablized secondary structure. Thus, we encoded compensatory adenine to cytosine mutations within the stem region (A48, A49 position) of the mod2 sgRNA variant and observed robust mutagenesis at the modified sgRNA locus (Fig. 1B). Additional variant sgRNAs (mod3, mod4 and mod5) did not exhibit noticeable self-targeting activity. Thus, the mod2 sgRNA was hereafter referred to and used as the stgRNA architecture.

We further characterized the mutagenesis pattern of the stgRNA by sequencing the DNA locus encoding it. A HEK 293T cell line expressing the stgRNA was transfected with a plasmid expressing either Cas9 (Construct 7, Table S2) or mYFP driven by the CMV promoter. Genomic DNA was harvested from the cells at either 24 hours or 96 hours post-transfection and subjected to targeted PCR amplification of the region encoding the stgRNAs. The PCR amplicons were either sequenced by MiSeq or cloned into *E. coli* for Sanger sequencing of individual bacterial colonies (Fig. S1). We found that cells transfected with the Cas9-expressing plasmid exhibited significant mutation frequencies in the stgRNA loci and those frequencies increased over time, compared to cells transfected with the control mYFP expressing plasmid (Fig. 1C). By using high-throughput sequencing, we inspected the mutated sequences generated by stgRNAs to determine the probability of insertions or deletions occurring at specific base pair positions. We calculated the percentage of those that contained insertions or deletions at each base pair position amongst all mutated sequences (Fig. 1D). We observed higher rates of deletions compared to insertions at each nucleotide position. Moreover, an elevated percentage of mutated sequences exhibited deletions consecutively spanning nucleotide positions 13-17 for this specific stgRNA (20nt-1). We carried out a more thorough analysis into the sequence evolution patterns of stgRNAs, as described later in Fig. 3.

**Fig. 3.**
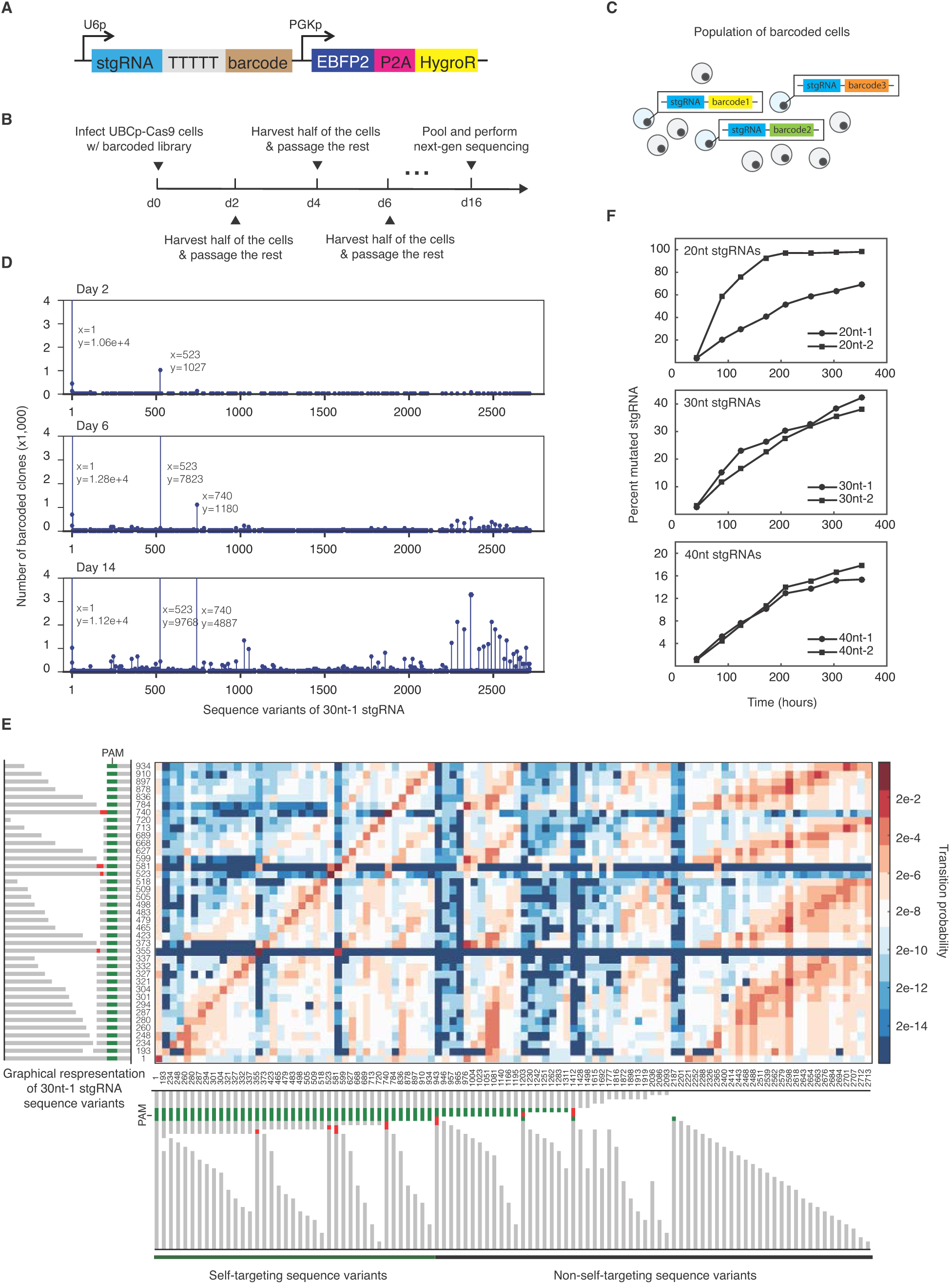
stgRNA sequence evolution analysis. (**A**) Schematic of the DNA construct used in building barcoded libraries encoding stgRNA loci. A randomized 16 bp barcode was placed immediately downstream of the stgRNA expression cassette in order to uniquely tag UBCp-Cas9 cells that contained integrated stgRNA loci. (**B**) The 16-day time course involved repeated sampling and passaging of cells in order to study sequence-evolution characteristics of stgRNA loci. (**C**) We lentivirally infected UBCp-Cas9 cells at an MOI ~0.3 so that the dominant population in infected cells contain single genomic copies of 16-bp-barcode-tagged stgRNA loci, which should be independently evolving. (**D**) The raw number of 16 bp barcodes that were associated with any particular 30nt-1 stgRNA sequence variant was plotted on the y-axis for three different time points (day 2, day 6 and day 14). Each unique, aligned sequence is identified by an integer index along the x-axis. The starting stgRNA sequence is shown as Index #1. (**E**) A transition probability matrix for the top 100 most frequent sequence variants of the 30nt-1 stgRNA. The color intensity at each (x, y) position in the matrix indicates the likelihood of the stgRNA sequence variant in each row (y) transitioning to an stgRNA sequence variant in each column (x) within the defined time scale (2 days). Since the non-self targeting sequence variants (which contain mutations in the PAM) do not participate in self-targeting action, the y-axis only consists of self-targeting stgRNA variants. The integer index of an stgRNA sequence variant is provided along with a graphical representation of the stgRNA sequence variant, wherein a deletion is illustrated using a blank space, an insertion using a red box, and an unmutated base pair using a gray box. The PAM is shown in green. From left to right on the x-axis and bottom to top on the y-axis, the sequence variants are arranged in order of decreasing distance between the mutated region and the PAM. When the distances are the same, the sequence variants are arranged in order of increasing number of deletions. (**F**) The percent mutated stgRNA metric is plotted for each of the stgRNAs as a function of time. We observe a reasonably linear range of performance metric for stgRNAs, especially for the longer SDS containing 30nt-1, 2 and 40nt-1, 2 stgRNAs. Also see Figs. S4-S7.

Given our observation that deletions are preferred over insertions, we suspected that stgRNAs would be shortened over time with repeated self-targeting activity, ultimately rendering them ineffective. To enable multiple cycles of self-targeting activity, we designed stgRNAs that are made up of longer SDSes. We initially built a cell line expressing an stgRNA containing a randomly chosen 30 nt SDS (Construct 8, Table S2) but did not detect noticeable self-targeting activity when the cell line was transfected with a plasmid expressing Cas9 (data not shown). We speculated that stgRNAs with longer than 20 nt SDSes might contain undesirable secondary structures that result in loss of activity. Therefore, we computationally designed stgRNAs that were predicted to maintain the scaffold fold of sgRNAs without undesirable secondary structures, such as stem loops and pseudoknots within the SDS (Methods). Stable cell lines encoding stgRNAs containing these computationally designed 30, 40 and 70 nt SDS (Constructs 9-11, Table S2) were transfected with a plasmid expressing Cas9 driven by the CMV promoter. T7 E1 assays of PCR amplified genomic DNA demonstrated robust indel formation in the respective stgRNA loci (Fig. 1E).

## stgRNA-encoding loci undergo multiple rounds of self-targeted mutagenesis

We sought to demonstrate that the stgRNA-encoding DNA locus in individual cells undergoes multiple rounds of self-targeted mutagenesis. To track genomic mutations in single cells over time, we developed a Mutation-Based Toggling Reporter (MBTR) system that generates distinct fluorescence outputs based on indel sizes at the stgRNA-encoding locus, which was inspired by a design previously described for tracking DNA mutagenesis outcomes (*11*). Downstream of a CMV promoter and a canonical ATG start codon, we embedded the Mutation Detection Region (MDR), which consists of a modified U6 promoter followed by an stgRNA locus. The MDR was immediately followed by out-of-frame green (GFP) and red (RFP) fluorescent proteins, which were separated by correspondingly out-of-frame ‘2A self-cleaving peptides’ (P2A and T2A) (Fig. 2A, Construct 13 in Table S2). Different reading frames are expected to be in-frame with the start codon depending on the size of indels in the MDR. In the starting state (reading frame 1, F1), no fluorescence is expected. In reading frame 2 (F2), which corresponds to any −1 bp frameshift mutation, an in-frame RFP is translated along with the T2A self-cleaving peptide, which enables release of the functional RFP from the upstream nonsense peptide. In reading frame 3 (F3), which corresponds to any −2 bp frameshift mutation, GFP is properly expressed downstream of an in-frame P2A and followed by a stop codon. We confirmed the functionality of this design by manually building constructs with stgRNA loci containing indels of various sizes (0 bp, −1 bp and −2 bp corresponding to Constructs 13, 14 and 15 in Table S2, respectively), and introducing them into cells without Cas9. We observed the expected correspondence between indel sizes and fluorescence output (Fig. S2).

**Fig. 2.**
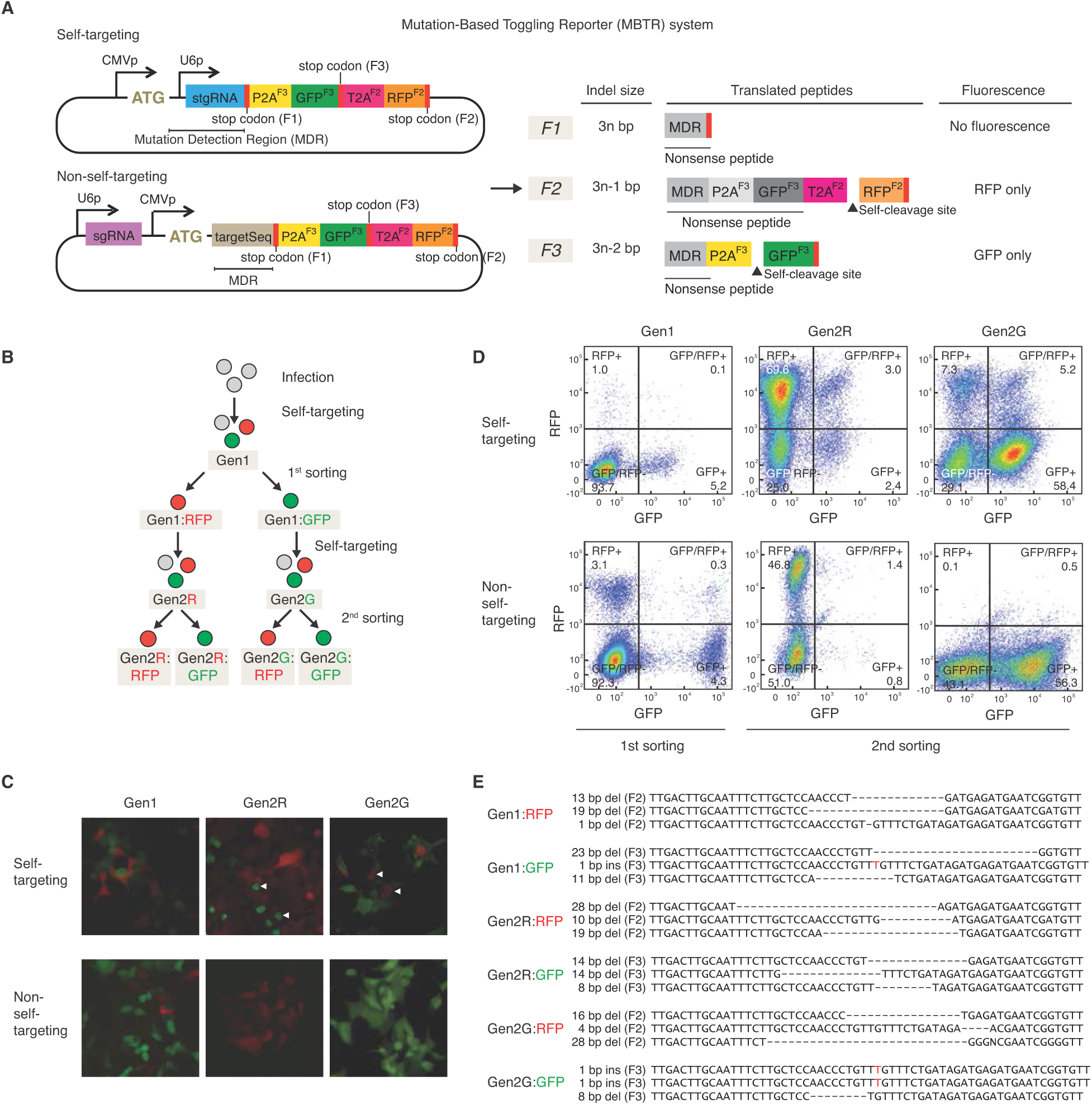
Tracking repetitive and continuous self-targeting activity at the stgRNA locus. (**A**) Schematic of the Mutation-Based Toggling Reporter system (MBTR system) consisting of an stgRNA in the Mutation Detection Region (MDR) or a regular sgRNA target sequence in the MDR region. We illustrate the expected fluorescent readouts of the MBTR system based on different indel sizes in the MDR. In the self-targeting scenario, a U6 promoter driven stgRNA with a 27 nt SDS is embedded between a constitutive human CMV promoter and modified GFP and RFP reporters. RNAP II mediated transcription starts upstream of the U6 promoter. Correct reading frames of each protein relative to the start codon are indicated in the superscript as F1, F2 and F3. Different sizes of indel formation at the stgRNA locus should result in different peptides sequences being translated. When translated in-frame, two ‘self-cleaving’ 2A peptides, P2A and T2A, are designed to cause co-translational ‘cleavage’ of the peptides and release functional fluorescent protein from the nonsense peptides, thus resulting in the appropriate fluorescent signal. The non-self-targeting construct consists of a U6 promoter driving expression of a regular sgRNA, which targets a sequence corresponding to the sgRNA embedded in the MBTR system as the MDR. (**B**) An outline illustrating the double-sorting experiment that tracks repetitive self-targeting activity using the MBTR system. HEK 293T cells stably expressing Cas9 (UBCp-Cas9 cells) were infected with MBTR constructs at low titre so that most of the infected cells had a single copy of the construct. Five days after the initial infection, generation 1 (Gen1) cells were sorted into RFP or GFP positive populations (Gen1:RFP and Gen1:GFP). The genomic DNA was extracted from a portion of the sorted cells. The rest of the sorted cells were allowed to grow to acquire further mutations at the stgRNA loci. The cells initially sorted for RFP or GFP fluorescence (Gen2R and Gen2G) were sorted again seven days after the first sort. The genomic DNA of the sorted cells (Gen2R:RFP, Gen2R:GFP, Gen2G:RFP and Gen2G:GFP) was collected, PCR amplified and Sanger sequenced after bacterial cloning. (**C**) Microscopy analysis and (**D**) flow cytometry data before the 1^st^ and 2^nd^ sorting of UBCp-Cas9 cells containing the self-targeting or non-self-targeting MBTR constructs. The white arrows in the microscope images indicate cells that expressed a fluorescent protein different from the one they were sorted for seven days earlier. For the self-targeting MBTR, we found that a significant fraction of cells expressed a fluorescent protein different from the one they were sorted for seven days earlier, indicating that repeated self-targeting mutagenesis that changed the indel sizes of the MDR (at least one mutagenesis event before the 1^st^ sort and at least one mutagenesis event before the 2^nd^ sort) occurred at each of the individual stgRNA loci present in the sorted cells. For the non-self-targeting MBTR, we observed minimal fractions of cells that expressed a fluorescent protein different from the one they were sorted for seven days earlier. (**E**) The genomic DNA collected from sorted cells was amplified and cloned into *E. coli*; the resulting bacterial colonies were then Sanger sequenced (Methods). A sample of Sanger sequences for the different sorted populations is presented along with their mutation type and the correct reading frame annotated. We observed a high correspondence between the mutated genotype and the observed fluorescent protein expression phenotype. Also see Fig. S2, S3.

We subsequently used the MBTR system to assess changes in fluorescent gene expression within cells constitutively expressing Cas9 to track repeated mutagenesis at the stgRNA locus over time. We built a self-targeting MBTR construct containing a computationally designed 27 nt stgRNA driven by a modified U6 promoter embedded in the MDR (Fig. 2A, Construct 13, Table S2). As a control, we built a non-self-targeting MBTR construct with a regular sgRNA that targets an identical 27 bp DNA sequence embedded in the MDR (Fig. 2A, Construct 16, Table S2). We integrated the self-targeting or the non-self-targeting construct (via lentiviral transduction at 0.3 multiplicity of infection (MOI) to ensure that most infected cells contained single copies) into the genome of clonally derived Cas9-expressing HEK 293T cells (hereafter called UBCp-Cas9 cells) and analyzed the cells by two rounds of Fluorescence-Activated Cell Sorting (FACS) based on RFP and GFP levels (Fig. 2B). In both cases, we found ~1-5% of the cells were RFP+ / GFP-or RFP-/ GFP+, which were sorted into Gen1:RFP and Gen1:GFP populations, respectively (Fig. 2C, D) and <0.3% cells expressed both GFP and RFP. We cultured the Gen1:RFP and Gen1:GFP cells for seven days, resulting in Gen2R and Gen2G populations, respectively. We then subjected the Gen2R and Gen2G populations to a 2^nd^ round of FACS. For cells with the stgRNA MBTR, a subpopulation of Gen2R cells toggled into being GFP positive, and similarly, a subpopulation of Gen2G cells toggled into being RFP positive. In contrast, cells containing the non-self-targeting MBTR with a regular sgRNA maintained their original fluorescence signals with no significant toggling behavior observed by FACS analysis (Fig. 2C, D). The toggling of fluorescence output observed in UBCp-Cas9 cells transduced with the stgRNA MBTR suggests that repeated mutagenesis resulting in multiple frameshifts in the MDR occurred at the stgRNA locus within single cells. To further corroborate this finding, we sequenced the stgRNA locus in individual cells from post-sorted populations in both rounds of sorting by cloning PCR amplicons into *E. coli* and performing Sanger sequencing on individual bacterial colonies (Fig. 2E and Fig. S3A). We found strong correlations (77%-100% accuracy) between the sequenced genotype and observed fluorescence phenotype in all of the sorted cell populations (Fig. S3B). Together, these results confirmed that repetitive mutagenesis can occur at the stgRNA locus within single cells.

## stgRNAs exhibit characteristic sequence evolution patterns

Having established that stgRNA loci are capable of undergoing multiple rounds of targeted mutagenesis, we set out to delineate their sequence evolution patterns over time. We hypothesized that we could infer characteristic properties associated with stgRNA sequence evolution by simultaneously investigating many independently evolving cell clones, all of which contain an exactly identical stgRNA sequence to start with (Fig. 3C). We synthesized barcoded plasmid DNA libraries in which the stgRNA sequence was maintained constant while a chemically randomized 16 bp barcode was placed immediately downstream of the stgRNA (Fig. 3A). Six separate DNA libraries were synthesized that encode stgRNAs containing six unique SDSes of different lengths: 20nt-1, 20nt-2, 30nt-1, 30nt-2, 40nt-1, or 40nt-2 (Constructs 19-24, Table S2). We used a constitutively expressed blue fluorescent protein, EBFP2, to confirm an MOI of ~0.3 so that most of the infected cells should contain single-copy integrants.

On day 0, lentiviral particles encoding each of the six stgRNA libraries were used to infect 200,000 UBCp-Cas9 cells in six separate wells of a 24-well plate. At a target MOI of 0.3, the infections resulted in ~60,000 successfully transduced cells per well. For each stgRNA library, eight cell samples were collected at time points approximately spaced 48 hours apart until day 16 (Fig. 3B). All samples from eight different time points across the six different libraries were pooled together and sequenced via Illumina NextSeq. After aligning the next-generation sequencing reads to reference DNA sequences (Methods), 16 bp barcodes that were observed across all the time points and the corresponding upstream stgRNA sequences were identified (Fig. S4A). For each of the stgRNA libraries, we found >10^4^ unique 16 bp barcoded loci that were observed across all of the eight time points (Fig. S4B). The aligned stgRNA sequence variants were represented with words composed of a four-letter alphabet: at each bp position, the stgRNA sequence was represented by either M, I, X or D, which stand for match, insertion, mismatch, or deletion, respectively (Fig. S4C, D, Methods). We identified >1000 unique sequence variants that were observed in any of the time points and any of the barcoded loci for each stgRNA (Fig. S5A, stgRNA sequences in Table S1). Although some sequence variants are found in common across the stgRNAs, majority of the sequence variants are unique to each stgRNA.

We plotted the number of barcoded loci associated with each unique sequence variant derived from the original 30nt-1 stgRNA for three different time points (Fig. 3D). Although the majority of the barcoded loci contained the original un-mutated stgRNA sequence (Index #1) for all three time points, we observed that a sequence variant containing an insertion at bp 29 (Index #523) and another sequence variant containing insertions at bps 29 and 30 (Index #740) gained significant representation by day 14. We noticed that most of the barcoded stgRNA loci evolved into just a few major sequence variants and thus sought to determine if these specific sequences would dominate across different experimental conditions. In Fig. S5B, we present the top seven most abundant sequence variants of the 30nt-1 stgRNA observed in three different experiments discussed in this work. The three experiments were performed with the 30nt-1 stgRNA encoded and: (1) tested *in vitro* in a HEK 293T-derived cell line (UBCp-Cas9), (2) tested *in vitro* in a HEK 293T-derived cell line in which Cas9 was regulated by the NF-κB-responsive promoter (inflammation-recording cells), or (3) tested *in vivo* in inflammation-recording cells (Fig. 3F, 4E and 4G, respectively). We found that six sequence variants (including Index #523 and #740) were represented in the top seven sequence variants for all three different experiments we performed with the 30nt-1 stgRNA. Moreover, even though we observed >1000 unique sequence variants for 30nt-1 stgRNA (Fig. S5A, stgRNA sequences in Table S1), these top seven most abundant sequence variants constituted >85% of the total sequences represented in each of these experiments. Thus, we speculate that stgRNA activity can result in specific and consistent mutations.

**Fig. 4.**
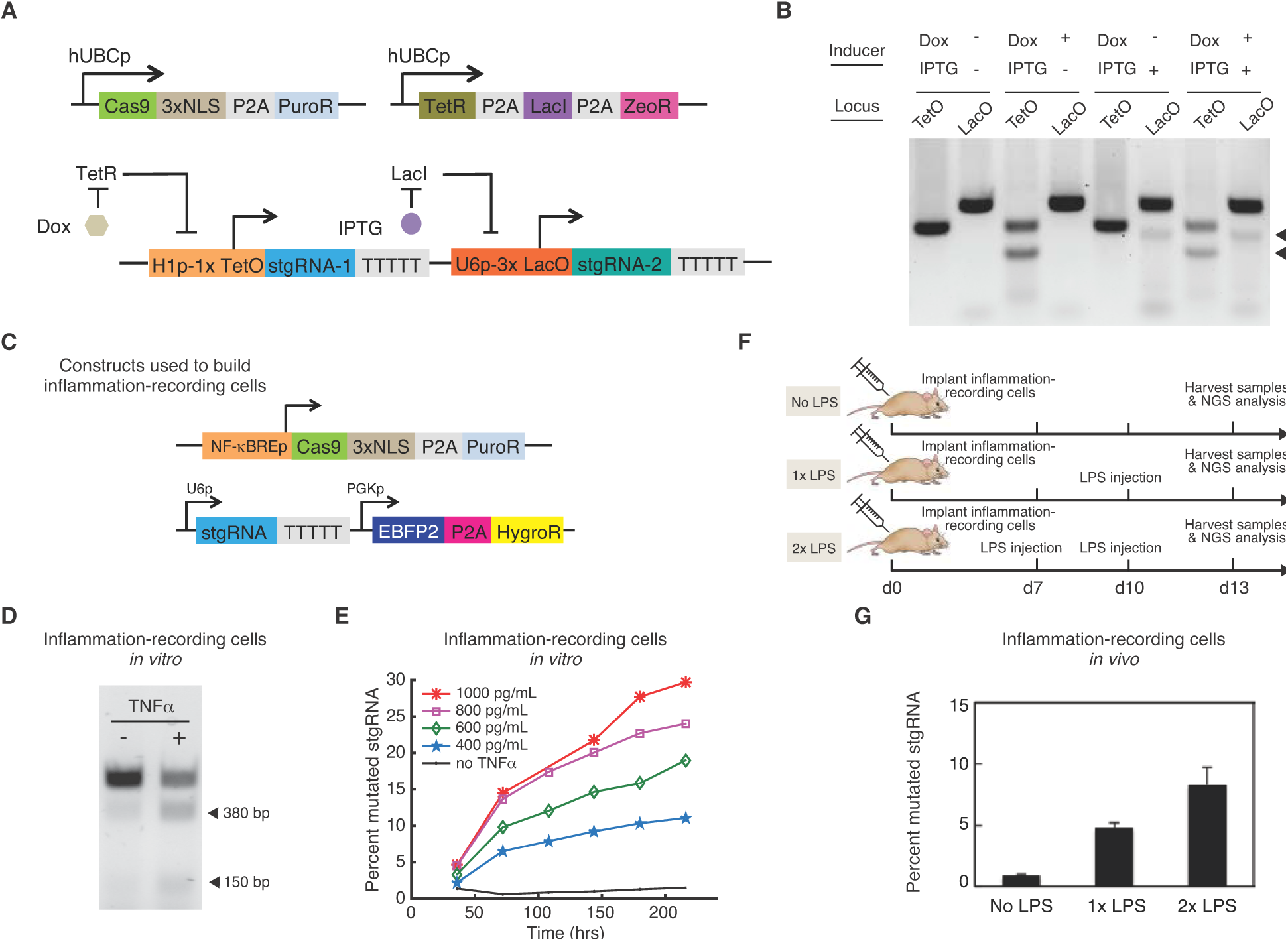
mSCRIBE as an analog memory device *in vitro* and *in vivo*. (**A**) Schematic of multiplexed doxycycline and IPTG-inducible stgRNA cassettes within human cell populations. By introducing small-molecule inducible stgRNA expression constructs into UBCp-Cas9 cells that also express TetR and LacI, the expression and self-targeting activity of each stgRNA can be independently regulated by doxycycline and IPTG, respectively. (**B**) mSCRIBE implements independently programmable, multiplexed genomic recording in human cells. Cleavage fragments observed from the T7 E1 assay of mSCRIBE units under independent regulation by doxycycline and IPTG are presented. Briefly, UBCp-Cas9 cells that express TetR and LacI were infected with lentiviral particles encoding the inducible stgRNA cassette from (**A**) and the cells were grown either in the presence or absence of 500 ng/mL doxycycline and/or 2mM IPTG. The cells were harvested 96 hrs post induction and PCR amplified genomic DNA was subject to T7 E1 assays. (**C**) Constructs used to build a HEK 293T-derived clonal NF-κBp-Cas9 cell line that expresses Cas9 in response to NF-κB activation. The 30nt-1 stgRNA construct was placed on a lentiviral backbone that expresses EBFP2 constitutively and was introduced in to NF-κBp-Cas9 cells via lentiviral infections at 0.3 MOI to build inflammation-recording cells. (**D**) T7 E1 assay testing for TNFα-inducible stgRNA activity in inflammation-recording cells *in vitro*. Inflammation-recording cells were grown either in the absence or presence of 1 ng/mL TNFα for 96 hrs. The genomic DNA was PCR amplified and assayed for the presence of mutations via the T7 E1 assay. We observed successful TNFα-inducible mSCRIBE activity in human cells. (**E**) Inflammation-recording cells were grown in media containing different amounts of TNFα or no TNFα. Cell samples were collected at 36 hr time point intervals for each of the concentrations. Genomic DNA from the samples was PCR amplified, sequenced via next-generation sequencing, and the percent mutated stgRNA metric was calculated. We observed graded increases in recording activity as a function of time and increasing concentrations of TNFα, thus demonstrating the analog nature of mSCRIBE. (**F**) Experimental outline for testing mSCRIBE in living mice. Inflammation-recording cells were implanted in the flank of three cohorts of four mice each. Three different cohorts of mice were treated with either no LPS, or with one or two dose(s) of LPS on days 7 and 10. After harvesting the samples on day 13 and PCR amplifying the genomic DNA followed by next-generation sequencing, the percent mutated stgRNA metric was calculated. (**G**) The percent mutated stgRNA metric calculated for the three cohorts of four mice is presented. The solid bars indicate the mean for each cohort (n = four mic in each condition) and the error bars indicate the s.e.m. mSCRIBE demonstrates increasing genomic recording activity with increasing doses of LPS in mice. Also see Figs. S8-10.

Given our observation that stgRNAs may have characteristic sequence evolution patterns, we sought to infer the likelihood of an stgRNA locus transitioning from any given sequence variant to another variant due to self-targeted mutagenesis. We computed such likelihoods in the form of a transition probability matrix, which captures the probability of a sequence variant transitioning to any sequence variant within a given time frame (Fig. 3E, Methods). Briefly, for every sequence variant observed in a future time point (daughter), a sequence variant from the immediately preceding time point and with the same barcode was chosen as the likely parent based on a minimal Hamming distance metric. Such parent-daughter associations were computed and normalized across all time points and barcodes to result in the transition probability matrix. In Fig. 3E, we present the transition probabilities of the top 100 most frequent sequence variants of the 30nt-1stgRNA. At each (x, y) position, we present the likelihood of a sequence variant from the y-axis (row) mutagenizing into a sequence variant on the x-axis (column) within a 2-day time frame. Since the presence of PAM is an absolute requirement for self-targeting activity, we only present the transition probabilities for sequences on the y-axis containing an intact PAM. From left to right on the x-axis and bottom to top on the y-axis in Fig. 3E, the sequence variants are arranged in order of decreasing distance between the mutated region and the PAM. When the distances are the same, the sequence variants are arranged in order of increasing deletions. We found that self-targeting sequence variants were generally more likely to remain unchanged than be mutagenized across the two-day time period, as indicated by high probabilities along the main diagonal (matrix elements where x=y) as annotated on Fig. S6. In addition, transition probability values were typically higher for sequence transitions below the main diagonal versus for those above the main diagonal, implying that sequence variants tend to progressively gain deletions (Fig. S6). Moreover, when compared with deletion-containing sequence variants, insertion-containing sequence variants tended to have a very narrow set of sequence variants they were likely to mutagenize into. Finally, we noticed that the predominant way in which mutated self-targeting sequence variants mutagenize into non-self targeting sequence variants is by losing the PAM and downstream region encoding the stgRNA handle while keeping the SDS encoding region intact.

Having analyzed the sequence evolution characteristics of stgRNAs, we envisioned that a metric could be computed based on the relative abundance of stgRNA sequence variants as a measure of stgRNA activity. Such a metric would enable the use of stgRNAs as intracellular recording devices in a population to store biologically relevant, time-dependent information that could be reliably interpreted after the events were recorded. From our analysis of stgRNA sequence evolution, we reasoned that novel self-targeting sequence variants at a given time point should have arisen from prior self-targeting sequence variants and not from non-self-targeting sequence variants. Thus, we calculated the percentage of sequences that contain mutations only in the SDS-encoding region amongst all the sequences that contain an intact PAM, which we call the percent mutated stgRNA, to serve as an indicator of stgRNA activity. In Fig. 3F, we plot the percent mutated stgRNA as a function of time for the six different stgRNAs. Except for the 20nt-2 stgRNA, which saturated to ~100% by 10 days, we observed non-saturating and steadily increasing responses of the metric for all stgRNAs over the entire 16-day experimentation period. Based on the rate of increase of the percent mutated stgRNA (percent mutated stgRNA / time), we note that stgRNAs encoding SDSes of longer length should have a greater capacity to maintain a steady increase in the recording metric for longer durations of time and thus should be more suitable for longer-term recording applications.

We also conducted a time course experiment with regular sgRNAs targeting a DNA target sequence to test their ability to serve as memory registers (Fig. S7). We used sgRNAs encoding the same 20nt-1, 30nt-2 and 40nt-1 SDSes tested in Fig. 3F (Constructs 25-27, Table S2) and found that unlike stgRNA loci, sgRNA target loci quickly saturate the percent mutated sequence metric and exhibit restricted linear ranges.

## Small-molecule inducible and multiplexed memory storage using mSCRIBE

We placed stgRNA loci under the control of small-molecule inducers to record chemical inputs into genomic memory registers. We designed doxycycline-inducible and isopropyl-β-D-thiogalactoside (IPTG)-inducible RNAP III promoters to express stgRNAs, similar to prior work with shRNAs (*12*, *13*) (Fig. 4A). We engineered the RNAP III H1 promoter to contain a Tet-operator, allowing for tight repression of promoter activity in the presence of the TetR protein, which can be rapidly and efficiently relieved by the addition of doxycycline (Construct 29, Table S2). Similarly, we built an IPTG-inducible stgRNA locus by introducing three LacO sites into the RNAP III U6 promoter so that LacI can repress transcription of the stgRNA, which is relieved by the addition of IPTG (Construct 30, Table S2). We first verified that doxycycline and IPTG-inducible stgRNAs worked independently when integrated into the genome of UBCp-Cas9 cells that also express TetR and LacI (Construct 28, Table S2) (Fig. S8). Next, we placed the doxycyline and IPTG-inducible stgRNA loci on to a single lentiviral backbone (Fig. 4A, Construct 31 in Table S2) and integrated them into the genome of UBCp-Cas9 cells that also expressed TetR and LacI. The induction of stgRNA expression by exposure to doxycycline or IPTG led to efficient self-targeting mutagenesis at the cognate loci as detected by the T7 E1 assay, while lack of exposure to doxycycline or IPTG did not (Fig. 4B). Moreover, when cells were exposed to both doxycycline and IPTG, we detected simultaneous mutation acquisition at both loci, thus demonstrating inducible and multiplexed molecular recording across the cell populations.

## Recording the activation of the NF-κB pathway via mSCRIBE

We next sought to build stgRNA memory units that record signaling events in cells within live animals. We adapted a well-established acute inflammation model involving repetitive intraperitoneal (i.p.) injection of lipopolysaccharide (LPS) in mice (*14*). Immune cells that sense LPS release tumor necrosis factor alpha (TNFα), which is a potent activator of the NF-κB pathway (*15*). The activation of the NF-κB pathway plays an important role in coordinating responses to inflammation (16). To sense the activation of the NF-κB pathway, we built a construct containing an NF-κB-responsive promoter driving the expression of the red fluorescent protein mKate2 (Construct 32, Table S2) and stably integrated it into HEK 293T cells. We observed a >50-fold increase in expression levels when these cells were exposed to TNFα *in vitro* (Fig. S9A, B, C). Next, we implanted these cells into the flanks of athymic nude mice (female nu/nu). After implanted cells reached a palpable volume, we performed i.p. injection of LPS and observed significant mKate2 expression (Fig. S9D) and elevated TNFα concentrations in the serum post LPS injection (Fig. S9E).

We then built a clonal HEK 293T cell line containing an NF-κB-induced Cas9 expression cassette (NF-κBp-Cas9 cells) and infected the cells with lentiviral particles encoding the 30nt-1 stgRNA at 0.3 MOI. These cells (hereafter referred to as inflammation-recording cells) accumulated stgRNA mutations, as detected with the T7 E1 assay, when induced with TNFα (Fig. 4D). We characterized the stgRNA memory unit in inflammation-recording cells by varying the concentration (within patho-physiologically relevant concentrations (*17*), Fig. S9E) and duration of exposure to TNFα *in vitro* and determining the percent mutated stgRNA metric (Fig. 4E). We observed graded increases in the percent mutated stgRNA metric as a function of time, thus demonstrating that stgRNA-based memory can record temporal information on signaling events in human cells. Furthermore, higher TNFα concentrations resulted in cells that had higher values for the percent mutated stgRNA metric, indicating that signal magnitude can modulate the mSCRIBE memory register in an analog fashion.

## Recording LPS-inducible inflammation *in vivo* via mSCRIBE

After characterizing the *in vitro* time and dosage sensitivity of our inflammation recording cells, we implanted them into mice. The implanted mice were split into three cohorts: no LPS injection over 13 days, an LPS injection on day 7, and an LPS injection on day 7 followed by another LPS injection on day 10 (Fig. 4F). The genomic DNA of implanted cells was extracted from all cohorts on day 13. The stgRNA locus was PCR amplified and sequenced via next-generation sequencing. We observed a direct correlation between the LPS dosage and the percent mutated stgRNA metric, with increasing numbers of LPS injections resulting in increased percent mutated stgRNA metric (also see Fig. S5B). Our results indicate that stgRNA memory registers can be used *in vivo* to record physiologically relevant biological signals in an analog fashion.

In Fig. 3E and 4F, we used PCR to amplify the stgRNA loci from ~30,000 cells and then calculated the percent mutated stgRNA metric as a readout of genomic memory. However, access to tissues or biological samples could be limited in certain *in* vivo contexts. To investigate the sensitivity of our stgRNA-encoded memory when the input biological material is restricted, we sampled 1:100 dilutions of the genomic DNA extracted from the TNFα-treated inflammation-recording cells in Fig. 4E (which corresponds to ~300 cells) in triplicate followed by PCR amplification, sequencing, and calculation of the percent mutated stgRNA metric (Fig. S10). We found very little deviation between the percent mutated sgRNA metric between samples with ~300 cells versus those from ~30,000 cells. We hypothesize that this tight correspondence is due to stgRNA evolution towards very few, dominating sequence variants, as was observed in Fig. 3D and Fig. S5B.

## Discussion and Conclusions

In this paper, we describe a novel architecture for self-targeting guide RNAs (stgRNAs) that can repeatedly direct Cas9 activity against the DNA loci that encode the stgRNAs. This technology enables the creation of self-contained genomic analog memory units in human cell populations. We show that stgRNAs can be engineered by introducing a PAM into the sgRNA sequence, and validate that mutations accumulate repeatedly in stgRNA-encoding loci over time with our MBTR system. After characterizing the sequence evolution dynamics of stgRNAs, we derive a computational metric that can be used to map the extent of stgRNA mutagenesis in a cell population to the duration or magnitude of the recorded input signal. Our results demonstrate that the percent mutated stgRNA metric increases with the magnitude and duration of input signals, thus resulting in long-lasting analog memory stored in the genomic DNA of human cell populations.

Because the stgRNA loci can be multiplexed for memory storage and function *in vivo*, this approach for analog memory in human cells could be used to map dynamical and combinatorial sets of gene regulatory events without the need for continuous cell imaging or destructive sampling. For example, cellular recorders could be used to monitor the spatiotemporal heterogeneity of molecular stimuli that cancer cells are exposed to within tumor microenvironments (*18*), such as exposure to hypoxia, pro-inflammatory cytokines, and other soluble factors. One could also track the extent to which specific signaling pathways are activated during disease progression or development, such as the mitogen-activated protein kinase (MAPK), Wnt, Sonic Hedgehog (SHH), and TGF-β regulated signaling pathways (*19*– *22*).

One limitation of our approach is that the NHEJ DNA repair mechanism is error-prone, so it is not easy to precisely control how each stgRNA cleavage event translates into a defined mutation, which could result in errors and noise in interpreting a given memory register. Ideally, each stgRNA cleavage event would result in a defined mutation, rather than a range of mutations. Amongst NHEJ repair mechanisms, recent studies have identified a more error-prone repair pathway, termed alternative NHEJ (aNHEJ). To enhance the controllability of mutations that arise over time, small molecule inhibitors of aNHEJ components, including ligase III and PARP1 could be used (*23*, *24*). The systematic engineering and characterization of a larger library of stgRNA sequences could also help to identify memory registers that are more efficient than the ones tested here.

Moreover, since our system generates a diverse set of stgRNA variants during the self-mutagenesis process, it is difficult to predict and eliminate potential off-target effects that may arise even if the original stgRNA can be designed for minimal off-target effects. As an alternative, we could fuse deactivated Cas9 (dCas9) to DNA cleavage domains such as single chain FokI nucleases (*25*) so that dCas9 could be targeted to a specific DNA locus with cleavage occurring away from the dCas9 binding site. This way, one can avoid generating new variants of stgRNAs that might target other sites in the genome while repeated targeting of the DNA locus can occur at locations distal to the dCas9 binding site, hence serving as a continuous memory register. Alternatively, adopting the recently described ‘base-editing’ strategy that employs cytidine deaminase (*26*) activity could help to avoid issues with using mutagenesis via DNA double strand breaks for memory storage. Epigenetic strategies, for example by fusing methyltransferases (*27*) or demethylases (*28*) to dCas9, could also be leveraged for continuous memory storage. Finally, in addition to recording information, this technology could be used for lineage tracing in the context of organogenesis. Embryonic stem cells containing stgRNAs could be allowed to develop into a whole organism and the resulting lineage relationships between multiple cell-types could be delineated via *in situ* RNA sequencing (*29*). In summary, we show that that mSCRIBE, enabled by self-targeting CRISPR-Cas, is useful for analog memory in mammalian cells. We anticipate that mSCRIBE will be applicable to a broad range of biological settings and should provide unique insights into signaling dynamics and regulatory events in cell populations within living animals.

## Acknowledgements

We thank members of the Lu lab for helpful discussions. We thank the MIT MicroBioCenter for technical support with next-generation sequencing and the MIT Koch Institute flow cytometry core facility for their technical assistance in cell sorting. This work was supported by the National Institutes of Health (DP2 OD008435, P50 GM098792), the Office of Naval Research (N00014-13-1-0424), the National Science Foundation (MCB-1350625) and NSF Expeditions in Computing Program Award #1522074. C.C. was supported by NSERC post-graduate fellowship.

### Author contributions

S.P., C.C., and T.K.L. conceived the work. S.P. and C.C. designed and performed experiments. S.P. performed computational analyses on next-generation sequencing data. C.C. conducted *in vivo* animal studies. S.P., C.C., and T.K.L. designed the experiments, interpreted and analyzed the data. S.P., C.C., and T.K.L. wrote the paper.

### Competing financial interests

T.K.L., S.P., and C.C. have filed a patent application based on this work with the US Patent and Trademark Office.

## Materials and Methods

### Vector construction

The vectors used in this study (Table S2) were constructed using standard molecular cloning techniques, including restriction enzyme digestion, ligation, PCR, and Gibson assembly. Custom oligonucleotides were purchased from Integrated DNA Technologies. The vector constructs were transformed into *E. coli* strain *DH5α*, and 50 μg/ml of carbenicillin (Teknova) was used to isolate colonies harboring the constructs. DNA was extracted and purified using Plasmid Mini or Midi Kits (Qiagen). Sequences of the vector constructs were verified with Genewiz and Quintara Bio’s DNA sequencing service. Sequences of all of the DNA constructs used in this work are listed in Table S2 and their plasmid maps are available at http://www.rle.mit.edu/sbg/resources/stgRNA/, password: stgRNA

### T7 Endonuclease I (T7 E1) assay and Sanger sequencing

Unless otherwise stated, cells used for T7 E1 assays were grown in 24-well plates with 200,000 cells per well. Genomic DNA from respective cell lines containing stgRNA or the sgRNA loci was extracted using the QuickExtract DNA extraction solution (Epicentre). Genomic PCR was performed using the KAPA-HiFi polymerase (KAPA biosystems) using the primers:

JP1710 – GCAGAGATCCAGTTTGGGGGGTTCCGCGCAC and JP1711 – CCCGGTAGAATTCCTCGACGTCTAATGCCAAC at 65°C for 30s and 25s/cycle extension at 72°C for 29 cycles. Purified PCR DNA was then used in the T7 Endonuclease I (T7 E1) assays. Specifically, 400 ng of PCR DNA was used per 20 μL T7 E1 reaction mixture (NEB Protocols, M0302). For Sanger sequencing, PCR amplicons from mutated genomic DNA were cloned in to KpnI/NheI sites of Construct 13 from previous work (*1*) and transformed into *E. coli* (DH5a, NEB). Single colonies of bacteria were Sanger sequenced using the Rolling Circle Amplification method (Genewiz, Inc).

### Cell culture, transfections and lentiviral infections

Cell culture and transfections were performed as described earlier (*1*). HEK 293T cells (ATCC CRL-11268) were purchased from and authenticated by ATCC. Our cell lines were tested negative for mycoplasma contamination by the Diagnostic Laboratory of the Division of Comparative Medicine at MIT. Lentiviruses were packaged using the FUGw backbone (*2*) (Addgene #25870) in HEK 293T cells. Filtered lentiviruses were used to infect respective cell lines in the presence of polybrene (8 μg/mL). Successful lentiviral integration was confirmed by using lentiviral plasmid constructs constitutively expressing fluorescent proteins or antibiotic resistance genes to serve as infection markers.

### Clonal cell lines and DNA constructs

A lentiviral plasmid construct expressing spCas9, codon optimized for expression in human cells fused to the puromycin resistance gene with a P2A linker was built from the taCas9 plasmid (*1*) (Construct 12, Table S2). The UBCp-Cas9 cell line was constructed by infecting early passage HEK 293T cells with high titre lentiviral particles encoding Construct 12 and selecting for clonal populations grown in the presence of puromycin (7 μg/mL). The NF-κBp-Cas9 cell line was built by infecting HEK 293T cells with high titer lentiviral particles encoding a NF-κB-responsive Cas9 expressing construct (Construct 33, Table S2). Transduced cells were induced with 1 ng/mL TNFα for three days followed by selection with 3 μg/mL puromycin. NF-κBp-Cas9 cells were then clonally isolated in the absence of TNFα. NF-κBp-Cas9 cells were infected with lentivirus particles encoding the 30nt-1 stgRNA locus at 0.3 multiplicity of infection (MOI) to build inflammation-recording cells. Cell lines used to test stgRNA activity were built by infecting HEK 293T cells with lentiviral particles encoding constructs 1 through 6 (Table S2) and selecting for successfully transduced cells with 300 μg/mL hygromycin. The cell line used to test inducible and multiplexed recording with doxycycline and IPTG was built by infecting UBCp-Cas9 cells with lentiviral particles encoding a DNA construct that expresses TetR and LacI constitutively (Construct 28, Table S2) followed by selection with 200 μg/mL zeocin for seven days.

### Flow cytometry and microscopy

Before analysis and sorting, cells were suspended in PBS with 2% fetal bovine serum. Cells were sorted using Beckmann Coulter MoFlo cell sorter. Flow cytometry analysis was performed with Becton Dickinson LSRFortessa and FlowJo. Fluorescence microscopy images of cells were obtained by using Thermo Scientific’s EVOS cell imager. The cells were directly imaged from tissue culture plates.

### Next-generation sequencing and alignment

Genomic DNA from respective cell lines was extracted using QuickExtract (Epicenter) and amplified using sequence specific primers containing Illumina adapter sequences P5 – AATGATACGGCGACCACCGAGATCTACAC and P7 – CAAGCAGAAGACGGCATACGAGAT as primer overhangs. Multiple PCR samples were multiplexed together and sequenced on a single flow cell using 8 bp multiplexing barcodes incorporated via reverse primers. The barcode library stgRNA samples in Fig. 3 were split into two groups and sequenced on the NextSeq platform (resulting in 154 and 178 million reads) while the 20nt-1 stgRNA samples in Fig. 1, the regular sgRNA samples in Fig. S7, TNFα dosage and time course characterization samples in Fig. 4E and the mouse tumor PCR samples in Fig. 4G were sequenced on the MiSeq platform (resulting in ~13 million reads per experiment). Paired end reads were assembled using the PEAR package (*3*). Optimal sequence alignment was performed by a custom written C++ code implementing the SS-2 algorithm (*4*) using affine gap costs with a gap opening penalty of 2.5 and a gap continuation penalty of 0.5 (see Code availability). The aligned sequences were represented using a four-letter alphabet in the ‘MIXD’ format where M represents a match, I represents an insertion, X represents a mismatch and D represents a deletion. At each base-pair position, the sequence aligned base pair is represented by one of the following letters: ‘M’, ‘I’, ‘X’ or ‘D’ (Fig. S4). 27 letter words were used to represent the 20nt stgRNA sequence variants wherein the 27 letters correspond to the first 20 bp of the SDS encoding region, followed by 3 bp of PAM and 4 bp representing the immediately adjacent 4 bp region encoding the stgRNA handle. Similarly, 37 and 47 letter words were used to represent the 30nt and 40nt stgRNA sequence variants.

### Barcoded stgRNA sequence evolution and transition probabilities

After sequence alignment, 16 bp barcodes and the stgRNA sequence variants (in the ‘MIXD’ format) were extracted. Only the 16 bp barcodes that were represented in all of the time points were considered for further analysis. All possible two-wise combinations of sequence variants associated with the same barcode but consecutive time points were evaluated for a ‘parent-daughter’ association. For every sequence variant in a future time point (a daughter), a sequence variant with the same barcode in the immediately preceding time point that had the minimum Hamming distance to the daughter sequence variant was assigned as the parent. Since the presence of an intact PAM is an absolute requirement for self-targeting capability of stgRNAs, only the sequence variants that contained an intact PAM were considered as potential parents. Many parent-daughter associations were computed across all the barcodes and time points, resulting in an overall count for each specific parent-daughter association. Finally, the counts were normalized such that the total likelihood of transitioning from each parent to all possible daughters would sum to one. The Hamming distance metric between two sequence variants in the ‘MIXD’ format was calculated by assigning a distance score for each base pair position. Specifically, if only one of the sequence variants being compared had an insertion at a particular base pair position, then the score for that position is assigned 2. In all other cases, the score at a base pair position was assigned 0 if the sequence variant letters were identical and 1 if they were not identical. The scores for each base pair position were summed up and used as the Hamming distance metric between the two sequence variants. Finally, while assigning parent-daughter associations, unless the parent and the daughter sequence variants were exactly identical, sequence variants that contain mutations in the PAM were not considered as potential parents.

### Design of longer stgRNAs

Longer stgRNAs were designed using the ViennaRNA package (*5*). Specifically, the RNAfold software was used to generate SDSes that retain the native structure of the guide RNA handle and no secondary structures in the SDS encoding region in the minimum free energy structure.

### *In vivo* inflammation model

Four to six weeks old female athymic nude mice (strain nu/nu) were obtained from the rodent breeding colony at Charles River Laboratory. They were specific pathogen free and maintained on sterilized water and animal food. All animals were maintained and used in accordance with the guidelines of the Institutional Animal Care and Use Committee. Sample sizes of the study were estimated based according to *in vivo* pilot studies and *in vitro* studies on the expected variance between animals and assay sensitivity. Inflammation-recording cells were suspended in matrigel (Corning, NY) in 1:1 ratio with cell growth media. 2 x10^6^ cells were implanted subcutaneously in the flank regions of mice. Animals were randomly assigned into experimental groups after tumor implantation with matched tumor sizes. Where indicated, mice were injected intraperitoneally with lipopolysaccharide (LPS) (from *Escherichia coli* serotype 0111:B4, prepared by from sterile ready-made solution from Sigma Chemical Co., St. Louis, MO) dissolved in 0.1 ml saline solution. Animal studies were conducted without blinding. The exclusion and inclusion criteria of the animal study were pre-established. Animals with tumors that grew more than 10 mm in its largest diameter during the experimental period were sacrificed and excluded from the study.

### Code availability

Relevant C++ routines used for data analysis can be found at:

http://www.rle.mit.edu/sbg/resources/stgRNA/ password: stgRNA

---

**Fig. S1.**
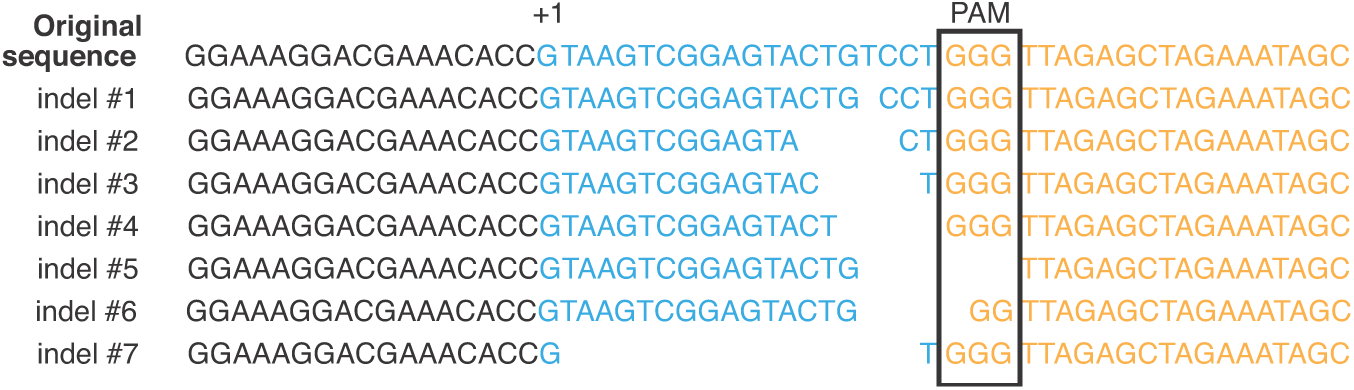
Sanger sequencing of stgRNA loci confirming self-targeting CRISPR-Cas activity. The stgRNA locus was PCR amplified from extracted genomic DNA. The purified PCR product was then digested by two restriction enzymes (KpnI/NheI) and cloned into a bacterial plasmid and transformed into *E. coli*. Bacterial colonies were picked next day and sequenced. The above indels were detected at the stgRNA loci. Also see Fig. 1C, D.

**Fig. S2.**
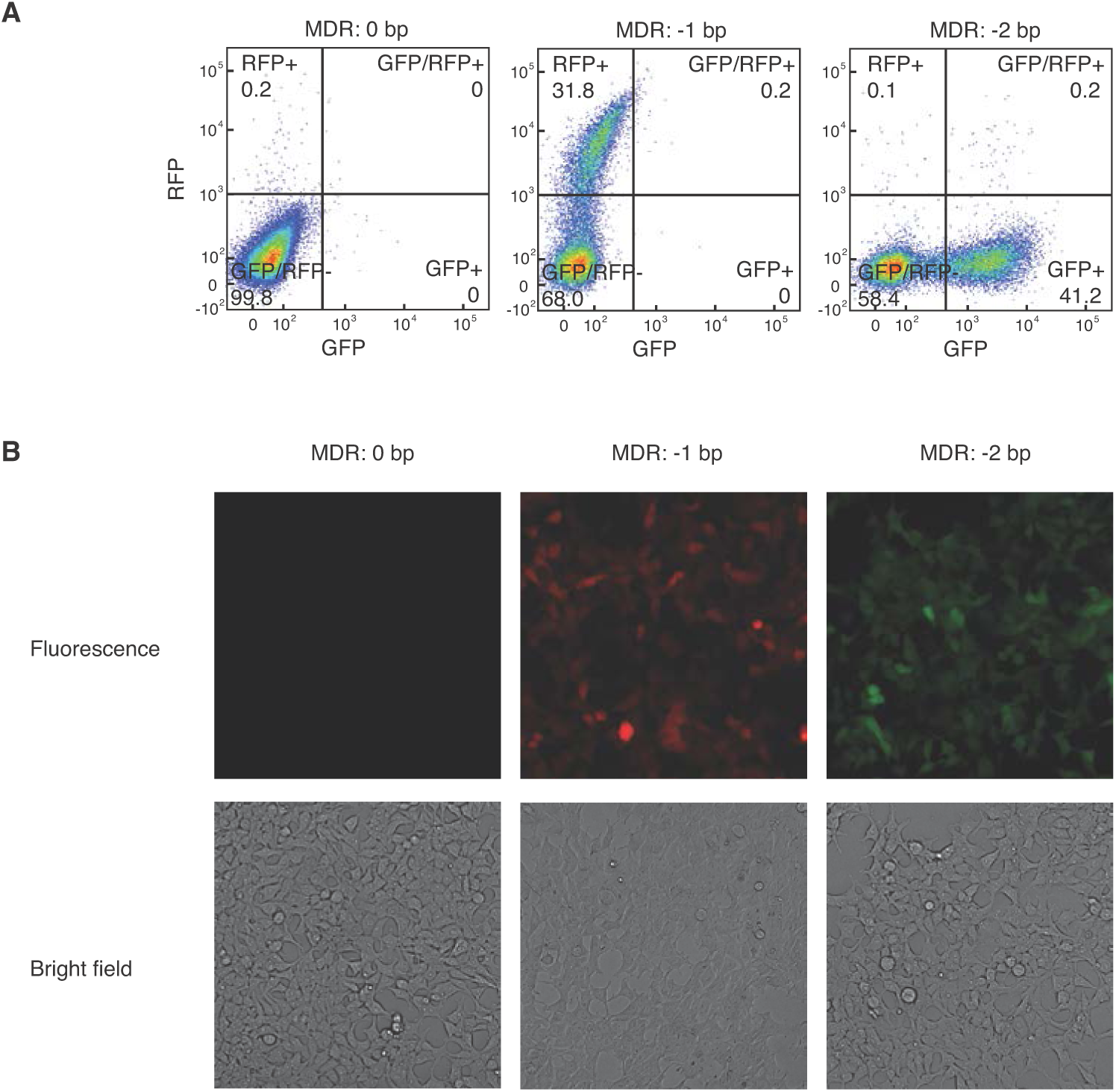
Validating functionality of the Mutation-Based Toggling Reporter (MBTR) system with different indel sizes in the Mutation Detection Region (MDR). We built MBTR constructs with stgRNAs containing indels of different sizes in the MDR (MDR: 0 bp = without an indel, MDR: −1 bp = with a −1 bp indel and MDR: −2 bp = with a −2 bp indel, Constructs 13-15, Table S2). We integrated these constructs into the genome of HEK 293T cells that do not express Cas9 via lentiviral infections at 0.3 MOI. We observed the expected correspondence between indel sizes in the MDR and fluorescence outputs as shown with flow cytometry analysis (top) and fluorescent microscopy (bottom). Also see Fig. 2A.

**Fig. S3.**
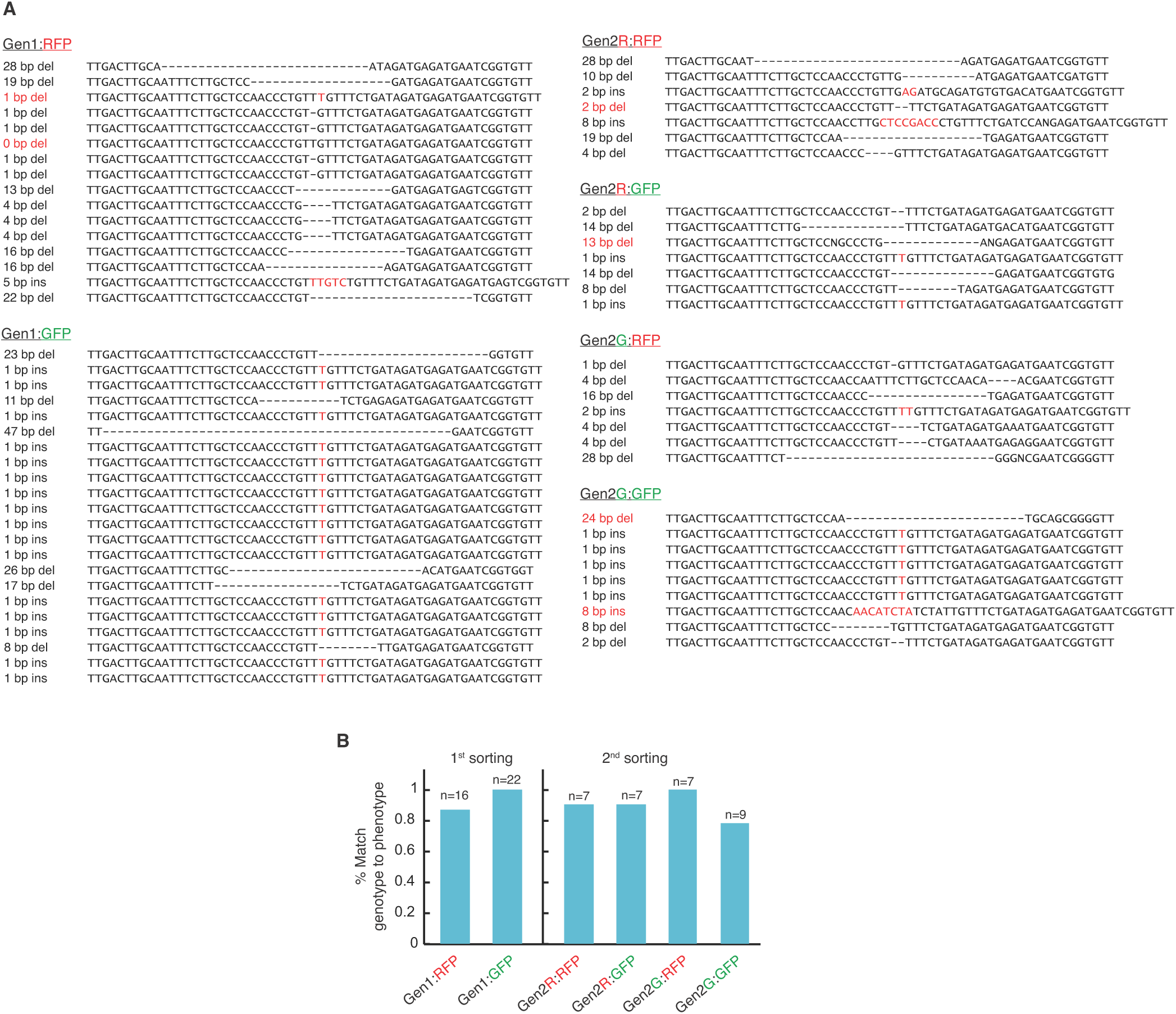
Sanger sequencing of stgRNA loci from sorted cells containing the self-targeting Mutation-Based Toggling Reporter (MBTR) construct. HEK 293T cells stably expresssing Cas9 (UBCp-Cas9 cells) were infected with lentiviral particles encoding the self-targeting MBTR construct at low titre. After 5 days, cells were sorted into RFP and GFP positive cells (Gen1:RFP and Gen1:GFP). The genomic DNA was extracted from half of the sorted cells, and stgRNA loci were amplified and cloned into *E. coli*. Individual bacterial colonies were then sequenced via Sanger sequencing (Methods). The other half of the sorted cells was allowed to grow and after a week from the initial sort, the cells were sorted again. The stgRNA loci of the harvested cells (Gen2R:RFP, Gen2R:GFP, Gen2G:RFP and Gen2G:GFP) were Sanger sequenced in a similar fashion. (**A**) Sanger sequencing data of each cell population. For each sequence, a description of the mutation is provided on the left annotated with reading frame (Fig. 2A). The descriptions indicated in red are mutations that do not correspond to the expected phenotype. We predominantly observed insertion sizes of only one or two bps but a wide range of deletion sizes. (**B**) We observed >80% match between the observed stgRNA sequence variant and the corresponding fluorescence phenotype in our samples. We speculate that the lack of perfect correspondence between the stgRNA sequence variant genotype and the fluorescence phenotype could be due to delays in the switching of cellular fluorescence following very recent self-targeted mutagenesis. Such delays could arise from the long half-lives of GFP and RFP (~24 hrs) and/or the time lag associated with gene expression. Also see Fig. 2.

**Fig. S4.**
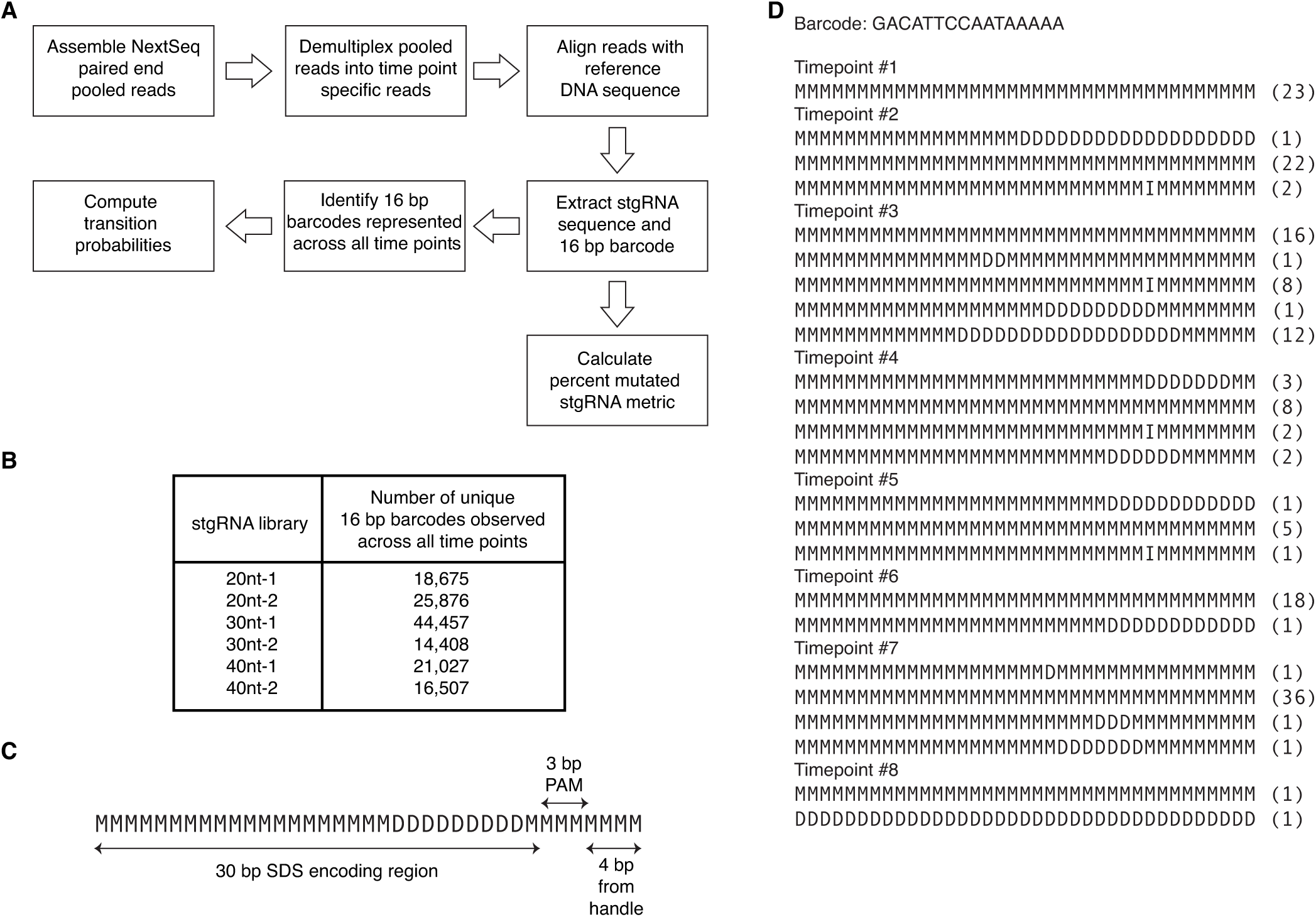
Computational analysis of stgRNA sequence evolution from the barcoded stgRNA library experiment. (**A**) Workflow illustrating the computational analysis employed in Fig. 3. Illumina NextSeq paired end reads for each of the six stgRNAs (20nt-1, 20nt-2, 30nt-1, 30nt-2, 40nt-1, 40nt-2) were assembled using PEAR (*3*). For each of the stgRNAs, assembled reads were binned into different time points after de-multiplexing using 8 bp indexing barcodes. Time-point-specific reads were then aligned with the reference DNA sequence (which is the sequence of the corresponding original, un-mutated stgRNA locus) using the SS2 affine-cost gap algorithm (*4*) implemented in C++ (Methods, Code availability). After aligning sequences with the reference, 16 bp barcodes and the potentially modified upstream stgRNA sequences were extracted. The aligned sequences were represented using words comprised of a four-letter alphabet in the ‘MIXD’ format where ‘M’ represents a match, ‘I’ an insertion, ‘X’ a mismatch and ‘D’ a deletion (Methods). For calculating transition probability matrices, “parent-daughter” associations that signify mutagenesis events that mutate a sequence variant from any given time point (parent) to a sequence variant (daughter) in the immediately following time point were computed. All possible pairs of sequences with the same barcode but from consecutive time points were evaluated for a potential parent-daughter association. For each daughter, a sequence variant with the same barcode from the immediate previous time point bearing the least Hamming distance was assigned a parent. A cumulative count of all parent-daughter associations was calculated across all barcodes and time points. Finally, to be a considered a true measure of probability, transition probabilities were normalized to sum to one. The “percent mutated stgRNA metric” was computed from the aligned sequences as the percentage of sequences that contain mutations in the SDS amongst all the sequences that contain an intact PAM. (**B**) The number of 16 bp barcodes represented across all the time points for each stgRNA. We observed >10,000 barcodes that were represented across all the time points for each stgRNA. (**C**) Example of an aligned sequence in the ‘MIXD’ format with base pair annotation. (**D**) Aligned sequences from a representative barcoded locus for the 30nt-1 stgRNA over time. For each barcode and each time point, unique sequence variants were identified. The parenthesis at the end of each of the sequence variants indicates the number of reads observed for that variant at each specific time point.

**Fig. S5.**
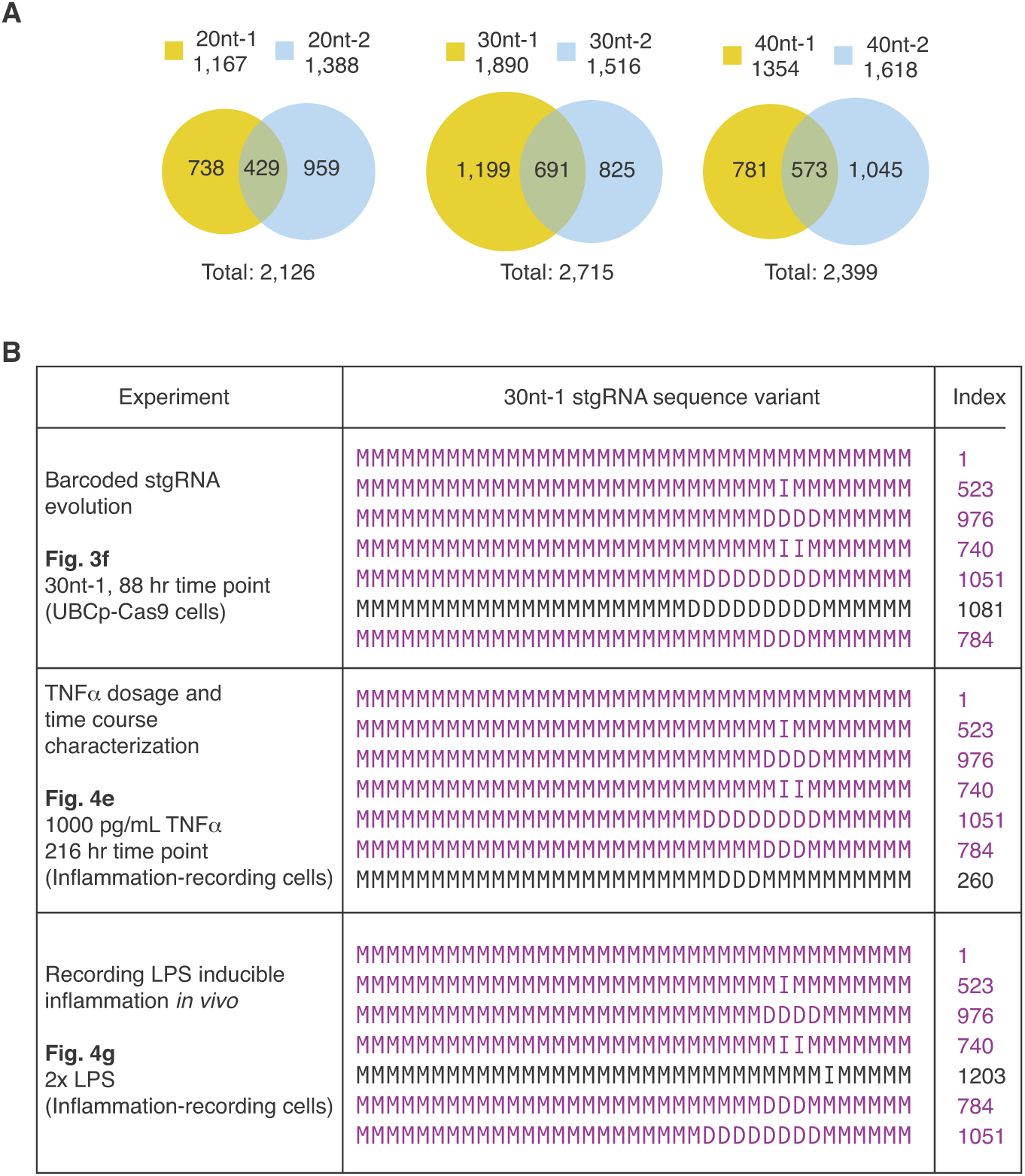
Analysis of stgRNA sequence variants in the ‘MIXD’ format. (**A**) Total numbers of stgRNA sequence variants in the ‘MIXD’ format observed for 20nt-1, 20nt-2, 30nt-1, 30nt-2, 40nt-1 and 40nt-2 stgRNAs in the barcoded stgRNA sequence evolution experiment across all time points. The numbers within the intersecting regions of the Venn diagrams are the number of sequence variants that are observed in common between 20nt-1 and 20nt-2, or 30nt-1 and 30nt-2, or 40nt-1 and 40nt-2 stgRNA loci. The numbers in the non-intersecting regions are the sequence variants observed exclusively with the respective stgRNA loci. Although some sequence variants are observed in common across stgRNAs, we noticed that a majority of stgRNA sequence variants are unique to each stgRNA. Also see Fig. 3D. (**B**) Top 7 most frequent 30nt-1 stgRNA sequence variants from three different experiments. After aligning the next-generation sequencing reads with the reference DNA sequence, sequence variants of the 30nt-1 stgRNA were extracted and represented in the ‘MIXD’ format. A 37 letter word is used to represent the 30nt-1 stgRNA sequence variants where the 37 letters correspond to the first 30 bp of the SDS encoding region, followed by 3 bp of PAM and a 4 bp of region from the stgRNA handle (Fig. S4C, D). The sequence variants presented above are the top 7 most frequently observed sequence variants of 30nt-1 stgRNA in three different experiments. The three experiments were performed with the 30nt-1 stgRNA encoded and: (1) tested *in vitro* in a HEK 293T-derived cell line (UBCp-Cas9), (2) tested *in vitro* in a HEK 293T-derived cell line in which Cas9 was regulated by the NF-κB-responsive promoter (inflammation-recording cells), or (3) tested *in vivo* in inflammation-recording cells respectively (Fig. 3F, 4E and 4G, respectively). Indices #1 through #2715 are assigned to denote each sequence variant of the 30nt-1 stgRNA. Six sequence variants highlighted in blue appear in the list of top 7 sequence variants across all three different experiments, implying that stgRNAs have consistent sequence evolution characteristics. Also see Fig. 3F, 4E and 4G.

**Fig. S6.**
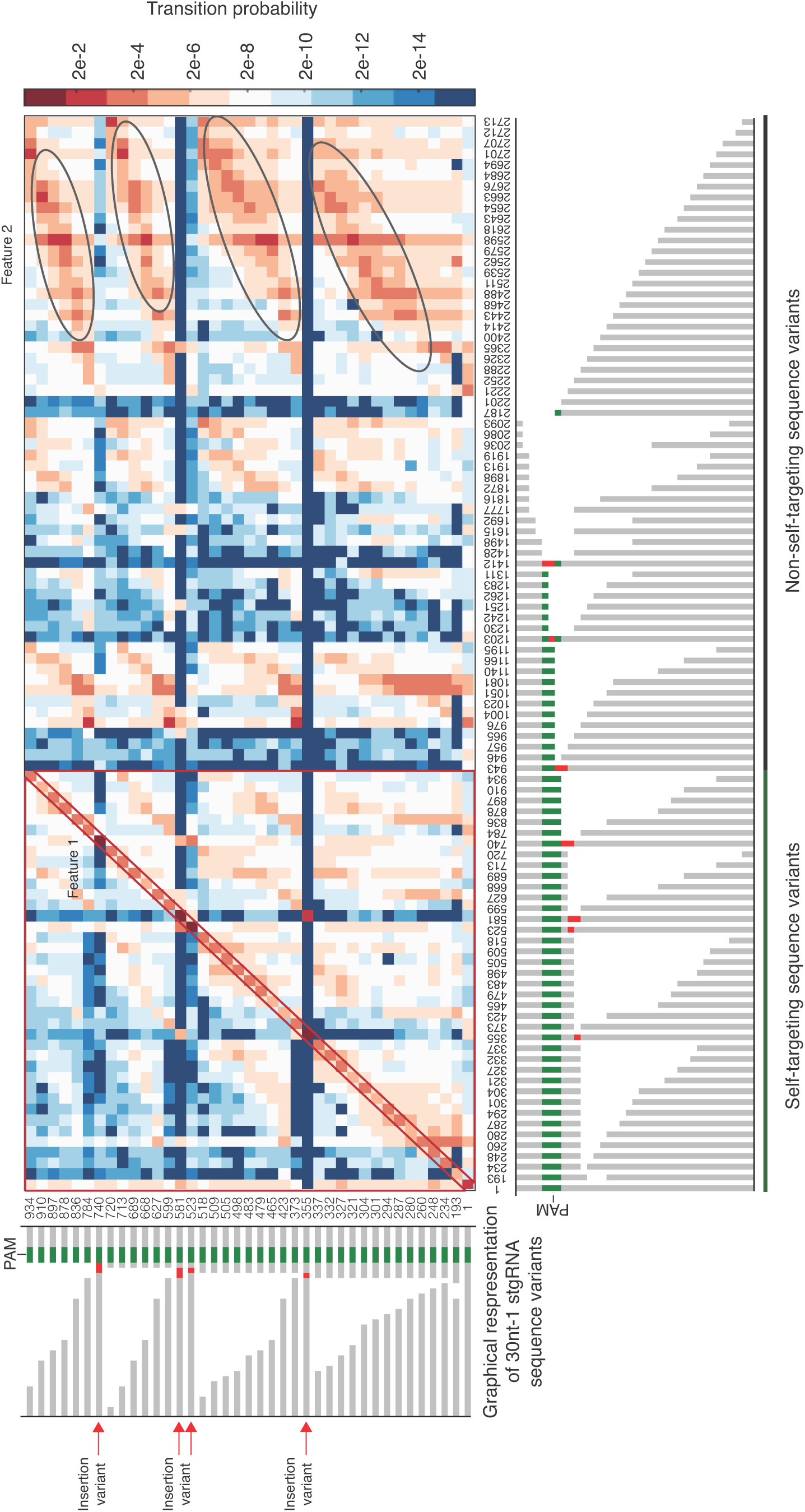
Transition probability matrix for 30nt-1 stgRNA. In the above plot, From left to right on the x-axis and bottom to top on the y-axis, the sequence variants are arranged in order of decreasing distance between the mutated region and the PAM. When the distances are the same, the sequence variants are arranged in order of increasing deletions. The highlighted features, Feature 1 and Feature 2, convey characteristic aspects of 30nt-1 stgRNA sequence evolution. In Feature 1, we observe that the transition probability values for transitions on the main diagonal (matrix elements that have x=y) are higher than those that are not on the main-diagonal, implying that the 30nt-1 stgRNA variants do not mutagenize much over a 48-hr time frame. We also observe that the transition probability values in the highlighted lower triangle (below the main-diagonal) are higher than the ones in the highlighted upper triangle (above the main-diagonal). This implies that 30nt-1 stgRNA sequence variants have a higher propensity to progressively gain deletions. In Feature 2, we observe that transition probability values are higher along the values indicated by the ovals. This implies that each of the mutated, self targeting stgRNA variants transition into non-self-targeting variants via mutagenic events that result in deletions of the downstream PAM sequences while keeping the upstream SDS-encoding regions intact. We also noticed that sequence variants containing insertions (highlighted by the red arrows) have a very narrow range of sequence variants they mutate into, compared with those that contain deletions. Also see Fig. 3E.

**Fig. S7.**
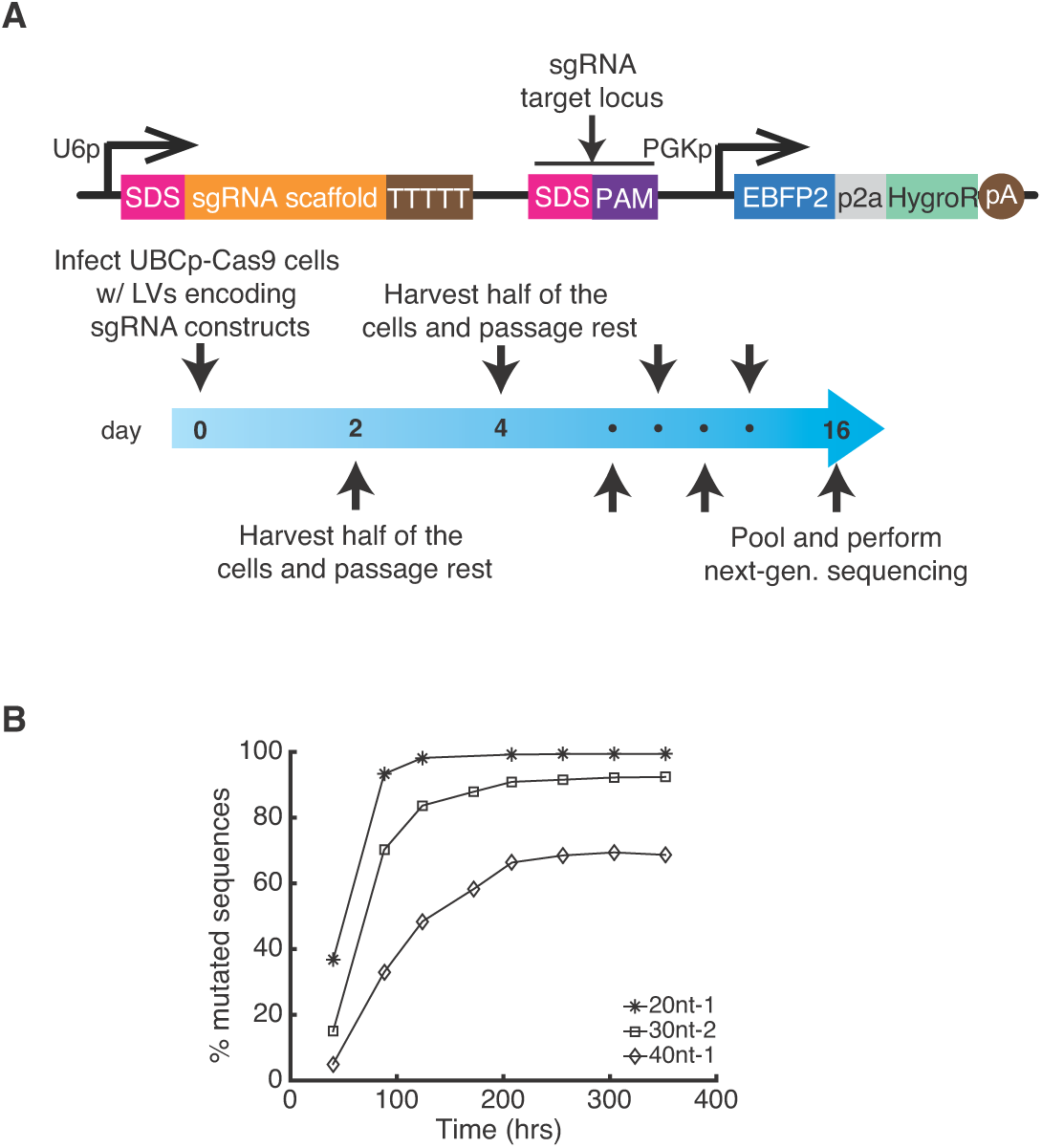
Regular sgRNAs as memory operators. (**A**) A circuit expressing a regular sgRNA that targets a DNA locus placed downstream was used in a time course experiment. The DNA constructs (Constructs 25-27, Table S2) are similar to the ones used for building the stgRNA barcode libraries in Fig. 3 A. The human U6 promoter drives expression of a regular sgRNA containing either a 20nt-1 or 30nt-2 or 40nt-1 SDS. An sgRNA target locus with its DNA sequence exactly homologous to the SDS and containing a downstream PAM (GGG, the identical PAM used in the stgRNA constructs) was placed 200 bp downstream of the RNAP III terminator ‘TTTTT’. The constructs encoding the 20nt-1, 30nt-2 and 40nt-1 sgRNAs were cloned into a lentiviral plasmid backbone harboring a constitutively expressed EBFP2, which is used an infection marker to ensure a target MOI of ~0.3. For each plasmid construct, ~200,000 UBCp-Cas9 cells were infected in separate wells of a 24 well plate on day 0 and cell samples were collected until day 16 at time points roughly spaced 48 hrs apart. At each time point, half of the cell population was harvested and the remaining half was passaged for processing at the next time point. All samples from eight different time points and three different SDSes were pooled together and sequenced in a high-throughput fashion via the MiSeq platform. After aligning each of the next-generation sequencing reads with the reference DNA sequences, potentially modified target loci were identified and mutation rates were calculated. (**B**) The percentage of target sequences mutated is presented as a function of time for the 20nt-1, 30nt-2 and 40nt-1 sgRNA designs. We found that conventional sgRNA-based memory units quickly saturate and do not exhibit long linear ranges when compared with stgRNAs. Thus, we concluded that conventional sgRNAs are not as amenable as stgRNAs for continuous recording over long time periods. Also see Fig. 3F.

**Fig. S8.**
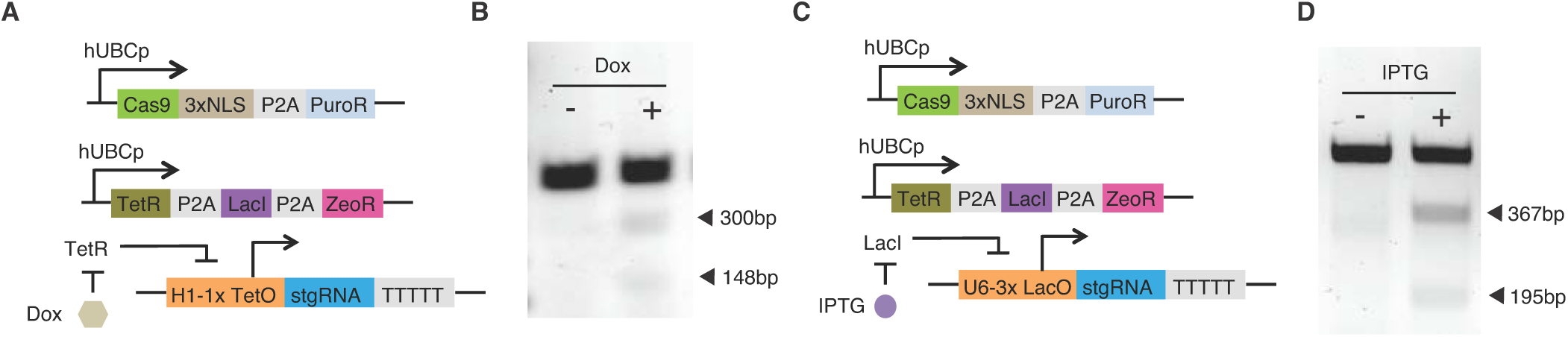
Small-molecule inducible mSCRIBE memory operators. Schematics of doxycycline (**A**) and IPTG (**C**) inducible stgRNA cassettes. By introducing small-molecule-inducible stgRNAs into UBCp-Cas9 cells also expressing TetR and LacI, stgRNA expression and its self-targeting activity can be tuned with the respective small molecules. (**A-B**) A doxycycline (Dox)-inducible stgRNA construct was built by introducing a Tet operator downstream of an H1 promoter (Construct 29, Table S2). The doxycycline-inducible stgRNA cassette was introduced into UBCp-Cas9 cells also expressing TetR and LacI. The cells were grown in the presence or absence of 500 ng/mL of doxycycline for 4 days and then assayed for self-targeted mutagenesis. The cleavage fragments observed from T7 E1 mutation detection assay showed that stgRNA activity was regulated by doxycycline. (**C-D**) Similarly, an IPTG-inducible stgRNA construct was built by introducing three copies of Lac operator within the U6 promoter (Construct 30, Table S2). The IPTG-inducible stgRNA cassette was introduced into UBCp-Cas9 cells also expressing TetR and LacI. The cells were grown in the presence or absence of 2 mM IPTG for 4 days and then assayed for self-targeted mutagenesis. In the presence of IPTG, mutations were detected in the stgRNA locus by the T7 E1 assay. Also see Fig. 4A, B and Constructs 28-31 in Table S2.

**Fig. S9.**
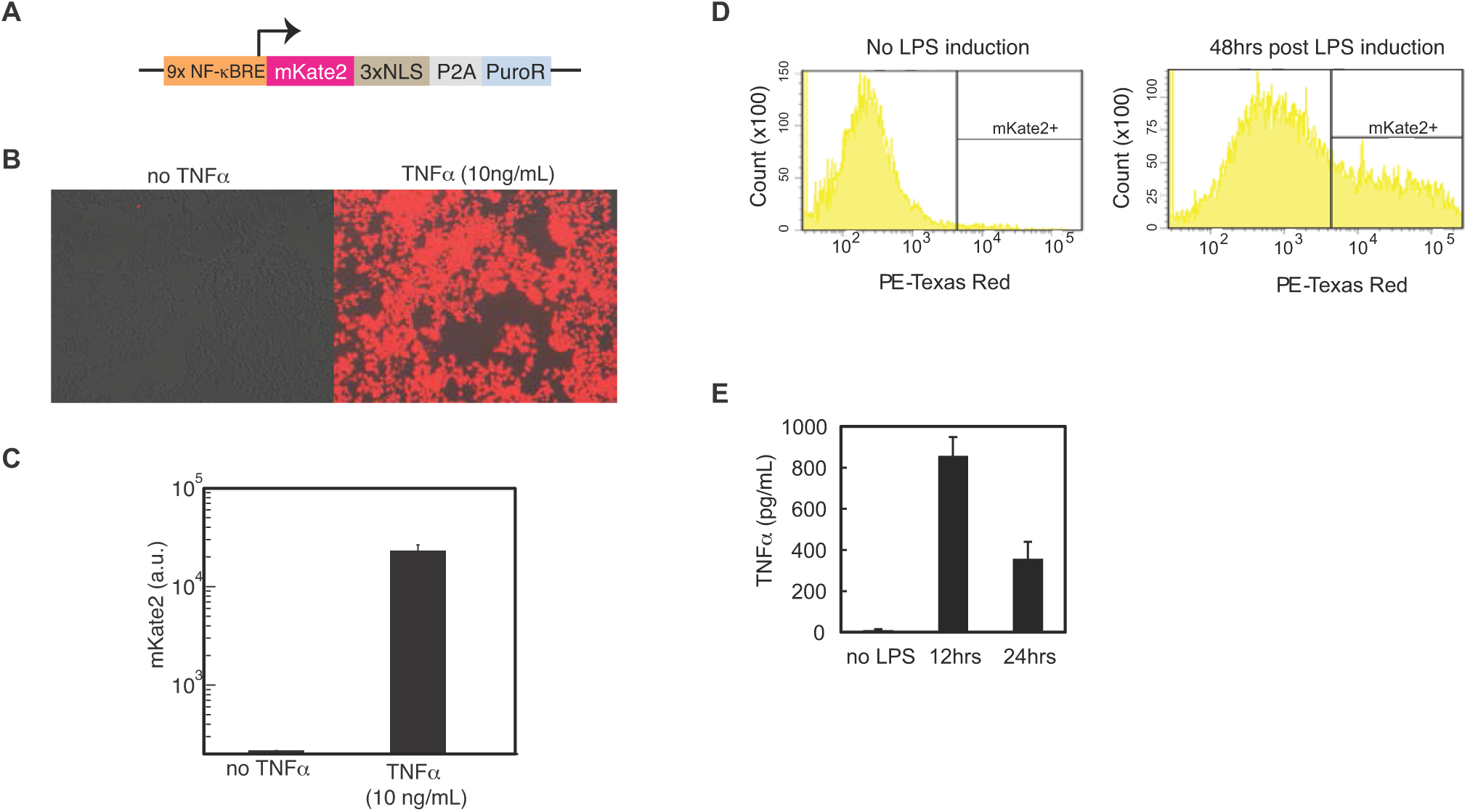
Characterization of *in vitro* (A-C) and *in vivo* (D-E) NF-κB-responsive gene expression to TNFα stimulation and LPS, respectively. (**A**) Schematic of an NF-κB-responsive mKate2 reporter vector construct. mKate2 expression in HEK 293T cell lines stably infected with an mKate2 construct regulated by an NF-κB-responsive promoter (Construct 32, Table S2) was quantified. (**B**) Fluorescence microscopy images of NF-κB-responsive stable cell lines infected with an mKate2 construct regulated by an NF-κB-responsive promoter grown in the absence or presence of 10 ng/mL TNFα. (**C**) Corresponding quantification of the fluorescence microscopy data. Three different biological replicates of the experiment were performed wherein 200,000 cells were grown in 24 well plates in the absence or presence of 10 ng/mL TNFα. The height of the bar represents the mean expression level of mKate2 and the error bars indicate the standard error of the mean for each condition. (**D**) Cells transduced with an NF-κB-responsive mKate2 reporter construct were implanted in mice. Cell samples collected 48 hours after i.p. LPS injection showed significant elevation of mKate2 expression compared to cell samples collected from mice that did not receive an LPS injection. (**E**) TNFα concentration in serum after LPS injection. After i.p. LPS injection, mice were sacrificed at different time points and blood was collected via cardiac puncture (n = 3 for each cohort). Serum TNFα concentrations were quantified with a mouse TNFα ELISA kit. The height of the bar represents the mean level of TNFα and the error bars indicate the standard error of the mean for each time point. Elevated TNFα levels were observed at 12 and 24 hours after LPS injection. Observed levels of TNFα in the serum were used as a guide to determine physiologically relevant concentrations of TNFα for the experiment in Fig. 4E.

**Fig. S10.**
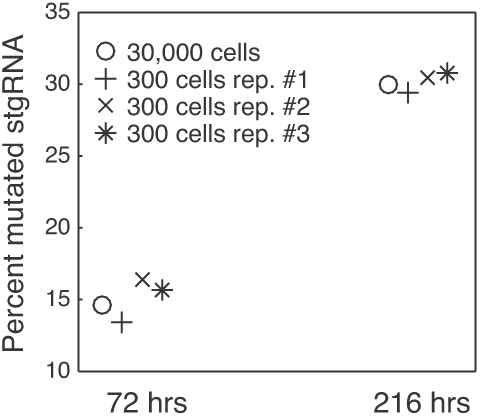
Percent mutated stgRNA metric calculated from sequencing genomic DNA corresponding to ~300 inflammation-recording cells, compared with the same metric measured from 30,000 inflammation-recording cells. Genomic DNA was harvested from inflammation-recording cells exposed to 1000 pg/mL TNFα in a 24-well plate. Half of the genomic DNA material (corresponding to ~30,000 cells) from the total genomic DNA per well (which was extracted from ~60,000 successfully infected cells) was PCR amplified and sequenced via next-generation sequencing. In addition, three samples of 1:100 dilutions of the genomic DNA obtained from the remaining half of the genomic DNA (corresponding to ~300 cells) were PCR amplified and sequenced via next-generation sequencing. The percent mutated stgRNA metric was calculated and plotted. We observed that as few as 300 cells can provide an accurate readout of the percent mutated stgRNA metric compared with 30,000 cells. Also see Fig. 4E.

